# Impact of DNA sequences in the DNA duplex opening by the Rad4/XPC nucleotide excision repair complex

**DOI:** 10.1101/2020.01.16.909549

**Authors:** Debamita Paul, Hong Mu, Qing Dai, Amirrasoul Tavakoli, Chuan He, Suse Broyde, Jung-Hyun Min

**Affiliations:** Department of Chemistry & Biochemistry, Baylor University, Waco, TX 76706, USA; Department of Biology, New York University, New York, NY 10003, USA; Department of Chemistry, University of Chicago, Chicago, IL 60637, USA

**Keywords:** Protein-DNA interactions, DNA damage recognition, DNA damage repair, nucleotide excision repair, sequence dependence, xeroderma pigmentosum, XPC, Rad4, X-ray crystallography, molecular dynamics simulations

## Abstract

Rad4/XPC is a key DNA damage sensor for nucleotide excision repair (NER) in eukaryotes. Rad4/XPC recognizes diverse bulky lesions by flipping out two lesion-containing nucleotide pairs and inserting a β-hairpin from the BHD3 domain (β-hairpin3) into the DNA duplex. We have previously observed that Rad4 can form the same ‘open’ structure when covalently tethered to a normal DNA sequence containing consecutive C/G’s (CCC/GGG) and that a similar open-like structure can be formed even when the β-hairpin3 is lacking. Here, we report a crystal structure of the Δβ-hairpin3 mutant tethered to a sequence containing alternating C/G’s (CGC/GCG). In contrast to the previous structures, Rad4 bound to CGC/GCG in a 180°-reversed manner, capping the end of the duplex without flipping out the nucleotides. MD simulations showed that CGC/GCG was inherently less ‘openable’ than CCC/GGG and that Rad4 failed to engage with its minor groove, a hallmark of productive binding towards ‘opening’. These results reveal that DNA sequences significantly influence the thermodynamic barrier for DNA opening by Rad4, which may render certain DNA structures/sequences resistant to ‘opening’ despite a long residence time of Rad4. The reverse-mode may indicate unproductive binding for NER whereas the DNA end-binding may hint at Rad4/XPC’s functions beyond NER.

## INTRODUCTION

DNA in the genome is under continuous attack from various environmental sources including ultraviolet (UV) radiation from sunlight and chemicals from food and pollutants (1, 2). The DNA lesions, if left unrepaired, can lead to mutations and to diseases such as cancer in humans (3). The nucleotide excision repair pathway is a versatile DNA repair mechanism that can remove the UV lesions and other bulky base adducts generated by chemical carcinogens (reviewed in (4, 5)). In transcription-coupled NER, the lesions are recognized by an RNA polymerase II obstructed by the adducts in the template strand during transcription. However, most of the DNA damage located elsewhere in the genome is recognized by the global-genome NER (GG-NER) which uses dedicated DNA damage sensors such as the XPC-RAD23B-Centrin2 protein complex. The XPC protein in particular is the subunit that specifically binds to and thus recognizes a wide variety of DNA lesions repaired by NER. The lesion-bound XPC complex in turn recruits the transcription factor IIH (TFIIH) containing the XPD helicase which unwinds and scans the DNA for damage verification. This in turn recruits XPA and replication protein A (RPA) that stabilizes the intermediate structure; subsequently the endonucleases (XPG and the XPF-ERCC1 complex) are summoned to incise the damage as a part of the 24-32 nucleotide-long single-stranded DNA. The resulting gap in the DNA is refilled by DNA repair synthesis and ligation. Mutations in the key NER proteins including XPC can lead to high susceptibility of cancers and neurological disorders as manifested by the xeroderma pigmentosum syndrome.

The key to the versatility of NER lies in its first lesion recognition step, and the mechanism by which XPC recognizes diverse NER lesions has been under extensive investigation. The crystal structures of the yeast XPC-RAD23B ortholog, Rad4-Rad23 (hereafter Rad4), bound to DNA lesions showed that the binding caused two nucleotide pairs harboring the lesion to be flipped out of the DNA duplex and a β-hairpin from the β-hairpin domain 3 (BHD3) was inserted into the DNA duplex to fill the gap (6, 7). Notably, in this ‘open’ structure, Rad4 did not directly contact the nucleotides harboring the lesion themselves (such as 6-4PP or CPD) but exclusively interacted with the nucleotides opposite the lesions. The structures therefore suggested that the protein must recognize various lesions in an indirect mechanism that relies on features of helix destabilization or distortion induced by a lesion rather than the lesion structure itself (8–11). Interestingly, we have also observed that Rad4 can form the same ‘open’ structure when covalently tethered to a non-specific, undamaged DNA sequence containing a stretch of CCC/GGG (12). This and other kinetic studies subsequently suggested that Rad4/XPC uses a kinetic-based lesion recognition mechanism where the protein may ‘open’ even an undamaged DNA site if given a sufficiently long residence time such as provided by the chemical tethering (‘kinetic gating’) (12–14). We also recently found that a similar open-like structure could be formed when Rad4 lacked the tip of β-hairpin2 or β-hairpin3 on the same DNA containing CCC/GGG (15). However, it is not yet known if Rad4 can open DNA of any structure/sequence given a sufficient residence time or if there is a threshold that would prevent it from ‘opening’ certain DNAs.

Under non-tethered conditions, the lesion binding specificity by Rad4/XPC and the repair rates for different lesions vary greatly depending on the lesion. For instance, among the UV-lesions, the more distorting and dynamic 6-4PP is much more efficiently recognized by Rad4/XPC and repaired than the less distorting CPD (7,16–18). Striking differences also exist among different DNA mismatches, which are useful *in vitro* substrates for Rad4/XPC even if they are not repaired by NER in cells: the inherently more dynamic and conformationally heterogeneous CCC/CCC mismatches are a better substrate than more homogeneous B-DNA-like TAT/TAT mismatches (13, 19). It is thus apparent that the Rad4-DNA specific binding must be influenced by multiple factors that affect the local structure and dynamics of the DNA. These factors include not only the type of a lesion but also the sequence context of a given DNA site as well as the protein’s residence time on the DNA. In principle, such mechanisms have a potential to allow the protein to act as more than a lesion-sensing protein. Interestingly, more and more studies report that XPC functions as a transcriptional coactivator that localizes to and functions around DNA in the absence of an apparent NER lesion (20–24). The detailed mechanistic insights into the protein’s action can thus be illuminating for these studies as well.

Here we report the structure of the β−hairpin 3 tip-deletion mutant (hereafter Δβ- hairpin3) tethered to a matched DNA sequence containing a string of alternating CG/GC repeats (hereafter ‘CGC/GCG’). In contrast to the ‘open(-like)’ structures of Rad4 bound to a DNA containing a string of consecutive C/G’s (‘CCC/GGG’), Rad4 bound to CGC/GCG in a 180 °- inverted fashion, preferring the BHD2/3 domains positioned near the duplex ends of the DNA while leaving the DNA duplex ‘closed’. Subsequent MD simulations comparing CCC/GGG versus CGC/GCG duplexes in the absence and presence of Rad4 show that CCC/GGG is likely to occupy conformational domains that are closer to the ‘open’ conformation than CGC/GCG even in the absence of the protein and is more likely to be distorted towards the ‘open’ structure when bound to Rad4. Altogether, our study illustrates the sensitivity of the Rad4-induced DNA opening to subtle differences in the DNA’s sequence-dependent structural and dynamical properties and that prolonged residence time may be a necessary but not sufficient condition for DNA opening. It also shows that Rad4/XPC has a propensity to bind to the duplex ends of the DNA in the absence of an ‘openable’ DNA site. Such mechanistic characteristics may also help elucidate the biological functions of Rad4/XPC beyond NER.

## MATERIALS AND METHODS

### Preparation of Rad4–Rad23 complexes

The intact (‘WT’) and the Δβ-hairpin3 Rad4-Rad23 complex constructs are as published previously (6,12–14). Rad4 in both constructs spanned residues 101–632 and contained all four domains involved in DNA binding (**Figure 1A**). The WT complex has been shown to exhibit the same DNA-binding characteristics as the full-length Rad4-Rad23 complex (6). The Δβ-hairpin3 mutant complex (construct name: <137>) lacked the tip of the long β−hairpin in the BHD3 domain of Rad4 (residues 599-605) in the context of the WT construct. For crystallization, the Δβ-hairpin3 mutant (construct name: <SC41B>) also harbored V131C/C132S mutations in Rad4 to introduce site-specific disulfide crosslinking with DNA as done before with CCC/GGG DNA (12, 15).

**Figure 1.**
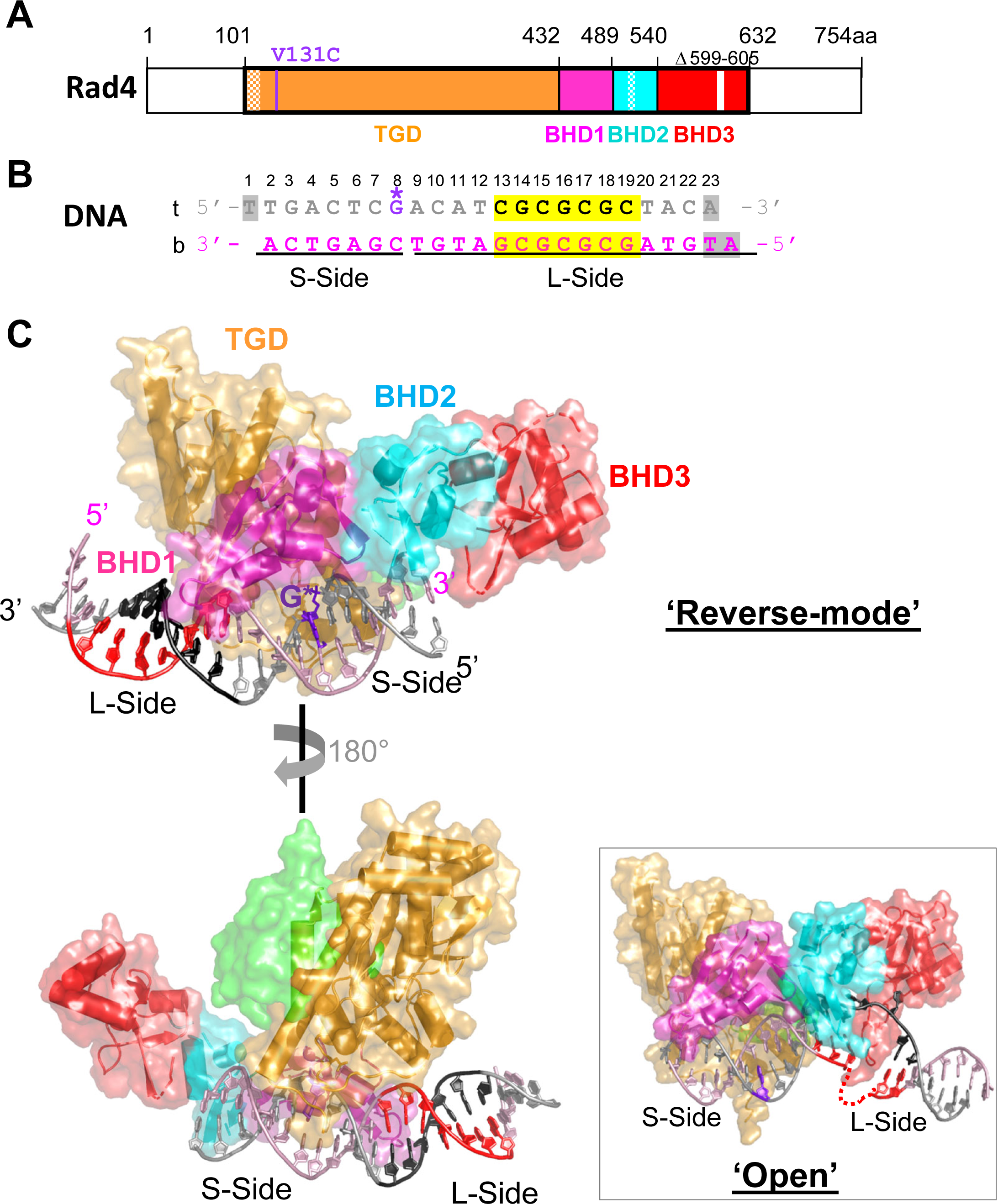
Crystal structure of the mutant Rad4–Rad23 complexes (Δβ-hairpin3) tethered to CGC/GCG matched DNA shows reverse-mode binding. (**A**) The crystallized Rad4 construct spans residues 101–632. The transglutaminase domain (TGD) is indicated in orange, β-hairpin domain 1 (BHD1) magenta, BHD2 cyan and BHD3 red. The deleted region in the BHD3 β- hairpin (residues 599-605) is indicated in white. The disordered regions in crystals (residues 101- 128, 518-525) are checkered. The V131C point mutation introduced for disulfide crosslinking is in purple. (**B**) The 23-bp CGC/GCG DNA construct for crystallization. Top strand (‘t’) is in silver and the bottom (‘b’) in pink. The disulfide-modified nucleotide, G* in dG8 is shown in purple. The CG repeats are highlighted in yellow and colored black in ‘t’ and red in ‘b’. The DNA residues with missing electron densities are shaded in gray. The bottom strand was the damage-containing strand in the ‘open’ structures of lesion-bound Rad4 (PDB ID: 2QSG, 6CFI) (6, 7). (**C**) The ‘reverse- mode’ structure of the Δβ-hairpin3 mutant bound to CGC/GCG DNA duplex (PDB ID: 6UG1). The color scheme is same as in (A) and (B). Rad23’s Rad4-binding domain (R4BD) is shown in light green. The bottom panel shows the structure rotated by 180 ° along a vertical axis. *(Inset)* The ‘open’ structure previously determined with CCC/GGG DNA tethered to the WT Rad4 complex (PDB ID: 4YIR).

All Rad4–Rad23 complexes in the study were co-expressed and purified from baculovirus-infected insect cells using previously described methods (6). Briefly, the Hi5 insect cells co-expressing Rad4 and Rad23 were harvested two days after infection. After lysis, the proteins were purified using immobilized metal affinity chromatography (Ni-NTA agarose, MCLAB) and then anion-exchange chromatography (Source Q, GE healthcare). For crystallization purposes, the complex was subjected to overnight thrombin digestion at 4 °C, followed by cation exchange (Source S, GE healthcare) and size-exclusion (Superdex200, GE healthcare) chromatography. The pure sample (> 90% by SDS PAGE) was concentrated by ultracentrifugation (Amicon Ultra-15, Millipore) to ∼15 mg/ml (185 μM) and stored in 5 mM bis-tris propane–HCl (BTP-HCl), 800 mM NaCl and 5 mM dithiothreitol (DTT), pH 6.8. For competitive EMSA studies, the protein complex was purified without thrombin digestion, thus retaining the UBL domain of Rad23 and a histidine-tag on Rad4 as also previously described (6, 12).

### Synthesis of oligonucleotides and preparation of duplex DNA

Different oligonucleotides containing disulfide-modified cytosine (the top strand) were prepared by incorporating the 2-F- dI-CE phosphoramidite (Glen Research) at the desired position during solid-phase synthesis. The conversion and deprotection of 2-F-dI were performed according to the guidelines provided by Glen Research (https://www.glenresearch.co/media/productattach/import/tbn/TB_2-F-dI.pdf).

Briefly, 2-F-dI-containing oligonucleotides were reacted with cystamine to tether the disulfide group, then deprotected with 1,8-diazabicycloundec-7-ene. All synthetic oligonucleotides were purified with HPLC. The HPLC purified bottom strands were purchased from IDT. Duplex DNA was prepared by annealing the top and bottom strands in ratio 1:1.1 in water.

### Disulfide crosslinking of the Δβ-hairpin3 mutant complex with double-stranded DNA

To crosslink the Δβ-hairpin3 mutant complex and DNA, dithiothreitol from the protein complex was first removed by extensive buffer exchange using a desalting column (Zeba Spin Desalting Column, 40,000 Da molecular weight cut-off, Thermo Scientific) pre-equilibrated in 5 mM BTP-HCl and 800 mM sodium chloride (NaCl), pH 6.8. The protein complex was thereafter incubated with DNA containing disulfide-modified base (1:1 molar ratio) in crosslinking buffer (5 mM BTP-HCl, 100 mM NaCl and 10% glycerol, pH 6.8) at 4°C overnight. The completion of the reaction was determined by SDS–PAGE under non-reducing conditions after treating the sample with 0.1 mM S-methylmethanethiosulfonate (Sigma) to quench the reaction. The crosslinking yield was ∼50–60%. The complex was then purified by anion-exchange chromatography (Mono Q, GE healthcare) over a 0–2 M NaCl gradient in 5 mM BTP-HCl and 10% glycerol, pH 6.8. The buffers were degassed by nitrogen purging. Purified crosslinked complex eluted at 400–480 mM NaCl and was further concentrated by ultrafiltration (Amicon, Millipore) to ∼3.5 mg/ml (30 μM).

### Crystallization of Δβ-hairpin3 mutant crosslinked with the CGC/GCG DNA duplex

To obtain crystals of the Δβ-hairpin3 complexed with matched DNA, we screened a panel of duplex DNA with varying lengths, sequences and overhang designs crosslinked to the mutant protein.

All crystallization trials were set up using the hanging-drop vapor diffusion method at 4 °C in which 1.5 μl of protein-DNA complex was mixed with 1.5 μl of crystallization buffer (50 mM BTP-HCl, 200 mM NaCl and 10–15% isopropanol pH 6.8) and sealed over 1 ml of crystallization buffer. Among the CGC/GCG DNA sequences of various lengths we tried (**Table S1**), the 22-bp DNA (CH9d) did not crystallize; the 24-bp (CH9a) and 25-bp (CH9b) DNA formed showers of needle-like microcrystals, but they did not diffract even after several rounds of optimization. The best crystals were obtained with the 23-bp DNA, first as showers of small plate-like crystals (20 μm) which appeared within a few days. Subsequently these crystals were harvested and used for micro-streak-seeding, which yielded larger crystals that grew to a maximum size of ∼70-80 μm in 10-12 days. The crystals were harvested in a harvest buffer (50 mM BTP-HCl, 200 mM NaCl, 3% isopropanol) and submerged for few seconds in a cryoprotectant buffer (50 mM BTP-HCl, 200 mM NaCl, 3% isopropanol and 20-25% MPD) before being flash-frozen in liquid nitrogen. Diffraction data were collected in LS-CAT, 21-ID-F beamline at 103 K and were processed with the HKL2000 (25). The data collection statistics are summarized in **Table S2**.

### Structure determination and refinement

The structure of the mutant Rad4–Rad23–DNA complex was determined by molecular replacement method using the previous structure of WT Rad4 crosslinked with CCC/GGG (PDB ID: 4YIR) using MOLREP (CCP4). Several rounds of model buildings were performed using WinCoot (26), followed by refinement with Phenix (27). First, the model building and refinement proceeded with the ‘open’ conformation as the model. However, the refinement statistics did not improve beyond R/Rfree: 33%/34% with this model.

Furthermore, the electron density of the DNA after the TGD-BHD1 domains appeared completely absent, indicating that the DNA may be bound in a different orientation. The reverse mode DNA model was constructed by 180° rotation of the DNA duplex with respect to the crosslinked nucleotide dG* (**Figure 1B**). This yielded improved model geometry and lower R/Rfree factor further confirming the validity of the alternate reverse mode model. The refinement statistics for the current model is summarized in **Table S2**. The final structure (PDB ID: 6UG1) contains residues 129–301, 305-504, 506-513, 528-598, 606-632 of Rad4 and 255– 311 of Rad23. The DNA residue numbers with missing densities are W1 and W23 in the top strand and Y1 and Y2 in the bottom strand. All figures were made using PyMOL Molecular Graphics System, version 2.1.1 (Schrodinger, LLC).

### Apparent binding affinities (Kd,app) determined by competition electrophoretic mobility shift assays (EMSA)

The specified Rad4-Rad23 complexes (0-300 nM) were mixed with 5 nM ^32^P-labelled DNA (matched or mismatched) in the presence of 1000 nM cold, undamaged DNA (CH7_NX) (**Figures S1**) in a binding assay buffer (5 mM BTP-HCl, 75 mM NaCl, 5 mM DTT, 5% glycerol, 0.74 mM 3-[(3-cholamidopropyl)dimethylammonio]-1-propanesulfonate (CHAPS), 500 µg/ml bovine serum albumin (BSA), pH 6.8). Mixed samples were incubated at room temperature for 20 min before being separated on 4.8% non-denaturing polyacrylamide gels. The gels were quantitated by autoradiography using Typhoon FLA 9000 imaging scanner (GE) and Image Lab software (Version 5.2.1 build 11, 2014; Bio-Rad). The apparent dissociation constants (Kd,app) were calculated as previously described using curve fitting by nonlinear regression using Origin software (OriginPro 9.6.0.172, Academic) (7,12,13,19).

### MD simulations and structural analyses

Full details of our MD simulation methods and analyses are given in the SI Methods.

## RESULTS

### Reverse-mode binding of Rad4 Δβ-hairpin3 mutant tethered to CGC/GCG matched DNA: Rad4 binds to the duplex end of the DNA instead of ‘opening’ the CGC/GCG sequence

In our previous study, we have shown that WT Rad4 formed an ‘open’ conformation with a matched DNA containing a central CCC/GGG sequence when covalently tethered to the DNA (12). We have also recently solved the structure of the β-hairpin3 tip deletion Rad4 mutant (residues 599-605; Δβ-hairpin3) tethered to the same CCC/GGG DNA, which showed a similar, yet more dynamical conformation (‘open-like’) (15). We have proposed that such ‘opening’ of a matched DNA was possible because the covalent tethering increased the protein’s residence time on a single register of DNA by limiting its diffusion: without tethering, a matched CCC/GGG DNA is a non-specific substrate for both the intact and the mutant Rad4 (6,12,13,19). However, it is unknown if such ‘opening’ would indeed happen for any sequence or structure given a long residence time (i.e., tethering) or if there is a selection barrier built into the protein’s DNA-opening mechanism that would only open select DNA sequences/structures even with a guaranteed long residence time.

To address these questions, we set out to determine the structures of the Rad4 tethered to DNA sequences other than the ‘openable’ CCC/GGG sequence. Since the Δβ-hairpin3 mutant was still able to form a ‘open-like’ structure with the CCC/GGG sequence like the WT under tethered conditions, we chose to focus on this mutant protein to make direct comparisons. After extensive trials, we obtained crystals with DNA duplexes containing a repeat of CG/GC dinucleotides (hereafter referred to as CGC/GCG). Like CCC/GGG, the CGC/GCG-containing duplexes also behave as non-specific substrates for Rad4 under non-tethered conditions in competitive gel-shift assays (**Figures S1**) and also has a comparable thermal stability as assessed by DNA duplex melting temperatures (**Figures S1 & S2**). The complexes with a 23-bp CGC/GCG DNA construct formed crystals in the *P1* space group (in contrast to those formed with the CCC/GGG DNA (*P41212*) that diffracted the best, up to 2.9 Å. The resulting structure was strikingly different from those previously determined with the CCC/GGG DNA: notably, the Rad4 was bound to DNA in a 180°-reversed orientation compared with the ‘open’ conformations formed with CCC/GGG. While the N-terminal transglutaminase domain (TGD) and BHD1 was still bound to the double-stranded portion of the DNA as in the ‘open’ conformation, the orientation of the protein with respect to the DNA was reversed by 180°. In such a mode of binding (hereafter referred to as ‘reverse mode’), the C-terminal BHD2/3 domains faced the short end of the DNA duplex with respect to the tethering site (hereafter referred to as ‘S-side’) instead of binding to and ‘opening’ the CGC/GCG site on the long end of the DNA (or ‘L-side’) (**Figure 1C**). The positioning of BHD2/3 on the ‘S-side’ also indicated that the BHDs would be largely blocking the duplex end of the DNA and that an extension of straight B-DNA would be incompatible with this binding conformation. On the other hand, TGD-BHD1 was bound to the ‘L-side’ of the DNA in which the CGC/GCG repeat sequence maintained its ‘closed’ duplex form. The DNA was also extended beyond TGD-BHD1 and made contacts with the BHD3 of Rad4 in a neighboring unit cell (**Figure S3**), which seemed more optimal for a 23-bp substrate than other DNA lengths. The structural parts common to the open(-like) and the reverse-mode structures superpose within ∼0.96 Å RMSD (**Figure S4**). However, the DNA in the reverse-mode structure maintained all base pairings without any nucleotide flipping or local unwinding seen in the ‘open’ structure (**Figure 2**).

**Figure 2.**
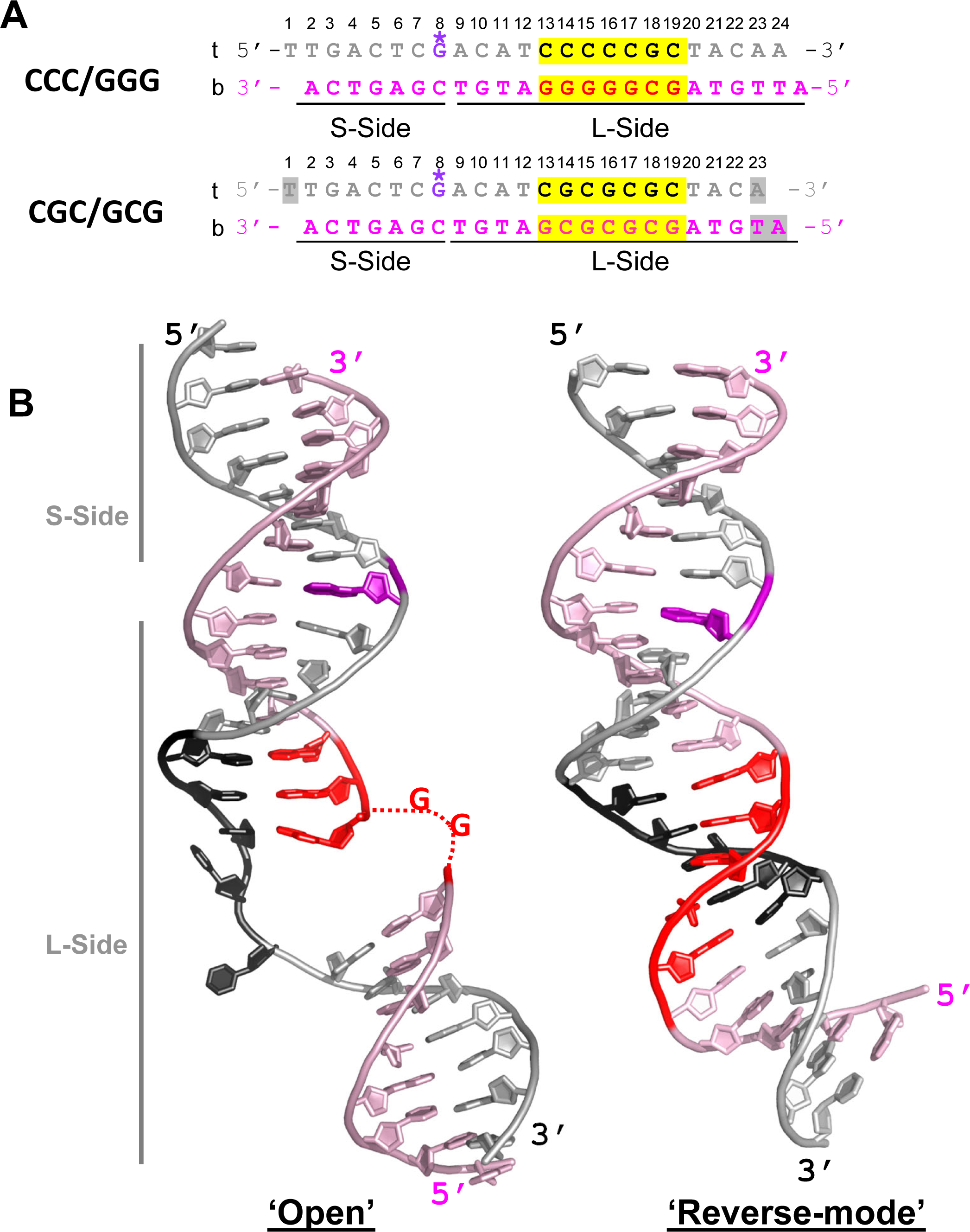
Comparison of DNA conformations in ‘open’ versus ‘reverse mode’ structures. **(A)** The **s**equences of the CCC/GGG DNA that formed an ‘open’ structure when tethered to the WT protein (PDB ID: 4YIR) and of the CGC/GCG DNA forming a ‘reverse-mode’ structure (PDB ID: 6UG1). The top (‘t’) and bottom strands (‘b’) are in silver and light pink respectively; CC/GG and CG/GC repeats are highlighted in yellow and colored black in ‘t’ and red in ‘b’. The DNA residues with missing electron densities are shaded in gray. The crosslinkable G*8 is in purple. The short (S-side) and long sides (L-side) of the DNA duplexes are designated with respect to the G* tethering site. **(B)** *(left)* The ‘open’ DNA conformation in the WT Rad4- CCC/GGG structure. *(right)* The ‘reverse-mode’ DNA shown in the Δβ-hairpin3-CGC/GCG DNA structure.

What causes Rad4 to bind to the CGC/GCG DNA in such a different binding mode from that with CCC/GGG in a complex whose constituents are otherwise the same? Although we cannot totally exclude the possibility that both ‘reverse’ and ‘open’ modes can co-exist in solution, it is reasonable to argue that the ‘reverse mode’ must be a preferred or dominant conformation for CGC/GCG as the ‘open’ mode is for CCC/GGG, and that this difference is mainly due to the DNA sequence difference rather than any other factors (see **SI Discussion**). The ‘reverse mode’ structure on CGC/GCG also supports the notion that not all DNA is ‘openable’ by Rad4 even given a long residence time provided by chemical tethering. To explore the structural factors that may influence the ‘open-ability’ by Rad4/XPC in detail, we have next turned to molecular dynamics (MD) simulations and examined the CCC/GGG and CGC/GCG DNA sequences without and with the bound Rad4. These studies show that the CCC/GGG and CGC/GCG sequences present different structures and dynamics even prior to Rad4 binding and demonstrate how their initial binding to Rad4 could be indicative of the distinct final structures observed in the crystals.

### MD simulations of unbound DNA duplexes reveal that the CCC/GGG duplex exhibits inherent structural distortions that foster the ‘open’ conformation

First, we investigated the impact of local sequence identity on the DNA conformations by performing 2 μs MD simulations for 13-mer DNA duplexes containing CCC/GGG and CGC/GCG sequences at their centers. Stable ensembles were achieved in the 0 – 2 μs range of production MDs following equilibration (**Figure S5**). Best representative structures are shown in **Figure 3A**. Our MD simulations of the unbound CCC/GGG and CGC/GCG 13-mer DNA duplexes revealed the striking impact of the run of guanines in CCC/GGG on the local DNA conformation. The structural ensembles of the CCC/GGG and CGC/GCG duplexes differ prominently in base pair slide, roll, twist, and also in helix bending direction (**Figures 3B and S6**). We first measured the 6 base pair step parameters: shift, slide, rise, tilt, roll and twist for the structures along the trajectories excluding the more dynamic end base pairs. Shift, rise and tilt did not show much deviation from an ideal B-DNA structure for either CCC/GGG or CGC/GCG sequences (**Figure S6A**). However, slide, roll, and twist did deviate from those of B-DNA and were different between the two sequences (**Figure 3B**). The CCC/GGG duplex had significant slide per GG step for the run of guanines with an average of ∼−1.5 Å, while the CGC/GCG duplex did not deviate much from the ideal B-DNA (0 Å slide) with an average slide of ∼−0.2 Å for the CG step and ∼−0.4 Å for the GC step. Correlated with its large slide, the CCC/GGG duplex exhibited consistent untwisting over the run of guanines indicated by an average twist angle per GG step of ∼30°, which is ∼6° lower than the ideal B-DNA value of 36° per step. In contrast, the CGC/GCG duplex showed only significant untwisting at its CG steps with an average twist angle of ∼31°, while the average twist angle for the GC steps is ∼35°. Correlated with the twist angle, CCC/GGG had a constant average roll angle per GG step of ∼8°, while CGC/GCG exhibited roll only at the CG steps with an average value of ∼9°. The base pair step parameters, twist, roll and slide, manifest the pair-wise sequence effects that have been well studied and explicated by Wilma Olson and colleagues (28, 29). Thus, the CG step and GC step alternate in the CGC/GCG sequence with less slide, lower twist, and greater roll at the CG than the GC step (**Figure 3B**). However, in the absence of such GC and CG alternations, the steric hindrance between guanine amino groups in the run of guanines have a dominant impact on the structure in the CCC/GGG sequence (28)

**Figure 3.**
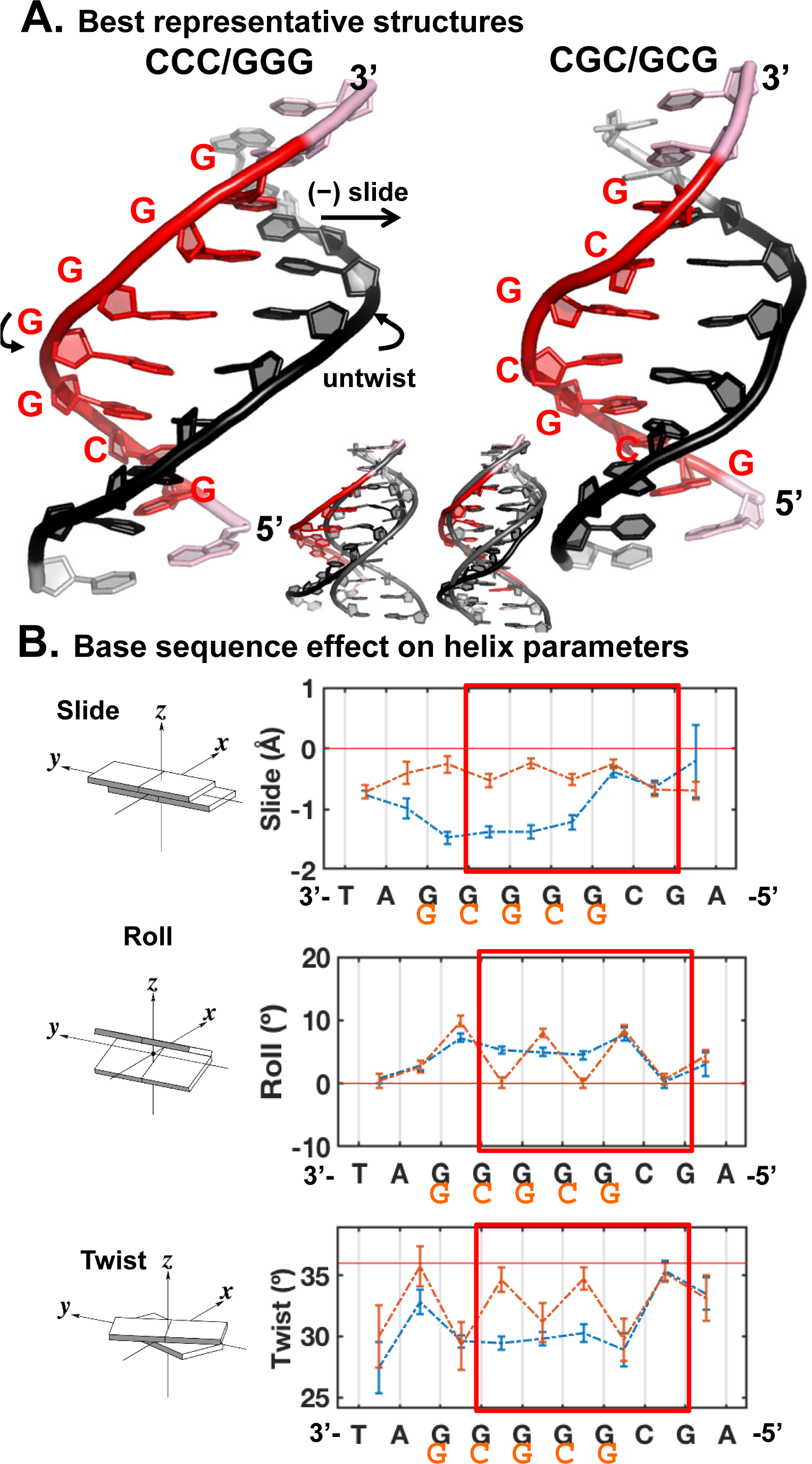
Intrinsic structural differences of the CCC/GGG and CGC/GCG sequences. (**A**) Best representative structures. **(B)** The DNA sequences had prominent impacts on the helix parameters, slide, roll and twist. See Figure S6 for the other parameters, shift, rise and tilt. Illustrations of the base pair step parameters are adapted from 3DNA (56). The block averaged means and standard deviations for the parameter values are shown. The twist angle is 36°/step for ideal B-DNA. Note that the sequence labels are from 3’ to 5’. The regions boxed red are centered around the putative ‘open’ site and the end base pairs of the 6-mer regions were used to calculate the (un)twist angle upon the DNA’s initial binding with Rad4 as shown in Figure 4.

We followed the dynamic bending of both duplexes by characterizing the DNA bend directions. While the bend angles themselves over the central 10-bp DNA were 13 ± 1° for both sequences (**Figure S6B**), the tail bend directions, calculated by using a pseudo-dihedral angle, were −45 ± 8° for CCC/GGG and −19 ± 10° for CGC/GCG (**Figure S6C**). This pseudo-dihedral angle adopts more negative values when bending is towards the minor groove around the potential open site. In the ‘open’ conformation crystal structure the bend direction dihedral is at −66°, while in the ‘reverse mode’ crystal of the Δbhd3-hairpin in complex with the CGC/GCG duplex it is −26°. Therefore, the directions of bending in the free, unbound CCC/GGG and CGC/GCG duplexes closely tracked with those of the respective DNA bound to Rad4 in the crystal structures.

Lastly, we have computed the van der Waals (vdW) energy for base stacking over the 6- mer region centered around the potential ‘open’ site (red box in **Figure 3B**). The CCC/GGG and CGC/GCG duplexes had a 1.3 kcal/mol difference over the central 6-mer region: −80.3 ± 0.3 kcal/mol for CCC/GGG vs. −81.6 ± 0.2 kcal/mol for CGC/GCG (Figure S6D).

In sum, MD simulations showed that CCC/GGG had higher slide, roll, and untwist compared to ideal B-DNA and more bending toward the ‘open’ conformation, accompanied by weaker van der Waals stacking energy compared with CGC/GCG. Such inherent distortions in free CCC/GGG DNA may lead to a higher propensity for the DNA to be ‘opened’ by Rad4 while CGC/GCG DNA could potentially resist such ‘opening’, which we further examined as described below.

### MD simulations of initial binding between Rad4 and the two different DNA sequences

Next, we asked how the DNA binding with Rad4 is directly impacted by the differences in the CCC/GGG and CGC/GCG sequences. For this purpose, we performed MD simulations on the initial binding process between the WT Rad4 and the two DNA sequences (see SI Methods). Prior MD simulation studies with different NER lesions have identified several features that are common to lesions repaired efficiently by NER: upon initial binding with Rad4, these lesion-containing duplexes all exhibit significant untwisting, ready engagement of the BHD2 b-hairpin (b-hairpin2) with the DNA minor groove, and capture of a partner base into a groove at the BHD2/BHD3 interface (7, 30). By contrast, NER-resistant lesions resisted such structural changes. In the current simulations, a stable BHD2 conformation in the minor groove was achieved at ∼1 μs for the CCC/GGG sequence, and for both simulations the conformations of the complexes were stable afterwards (**Figure S7**). Hence, we took the 1 – 2 μs trajectories as the initial binding states for further characterization. The best representative structures of each ensemble are shown in **Figure 4A**.

**Figure 4.**
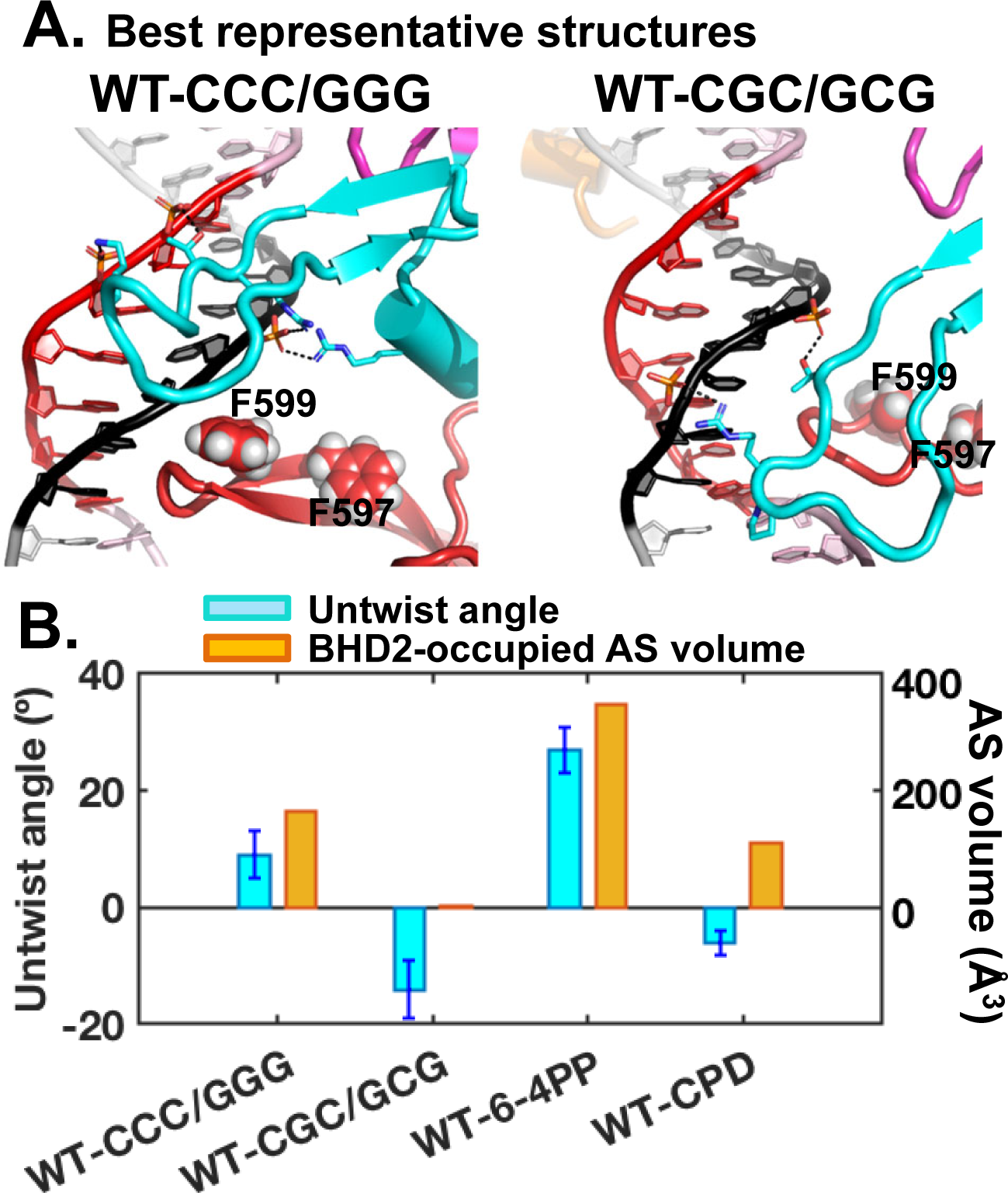
Initial binding of Rad4 to the CCC/GGG and CGC/GCG sequence-containing duplexes. **(A)** Best representative structures upon initial binding of Rad4. The structure is shown in cartoon representation color-coded as in Figure 1. Heavy atoms of the BHD2 amino acid side chains and the potential open site’s DNA backbone phosphate groups that form hydrogen bonds are shown in sticks. Hydrogen bonds are shown as black dashed lines. Side chains of two Phe (F599 and F597) in the β-hairpin3 are shown in spheres. **(B)** Potential ‘open’ site untwisting (6- mer) and BHD2 binding into the minor groove were quantified using the untwist angles and the BHD2-occupied AS volumes in the minor groove for the Rad4 binding. The values for the 6-4PP and CPD are from (7). The CCC/GGG sequence shows significant BHD2 binding and modest untwisting, resembling the well-recognized and repaired 6-4PP. In contrast, the CGC/GCG sequence shows no BHD2 binding and slight over-twisting, reminiscent of the poorly recognized/repaired CPD.

First, we computed the untwist angle of the duplex DNA around the potential ‘open’ site (see SI Methods, **Figure S8A)** (7, 30). The twist angles between the two base pairs that are 3-bp away from either side of the putative ‘open’ site (end base pair of the 6-mer sequence, boxed red in **Figure 3B**) were computed along the trajectories; the change in the twist angle from that of the initial twist angle was used as a measure of untwist around the ‘opening’ site: Untwist = Twist initial –Twist. The untwist angle thus reflects the extent of untwisting upon achieving the initial binding state. Positive values indicate untwisting and negative values indicate over-twisting. Upon achieving the initial binding state, the CCC/GGG duplex showed modest untwist of 9 ± 4°, whereas the CGC/GCG duplex showed over-twisting of −14 ± 5°. The positive value with CCC/GGG thus tracks with our previous result with the NER-proficient 6-4 photoproduct that showed 27 ± 4° of untwisting upon initial Rad4-binding; the value with CGC/GCG, on the other hand, is reminiscent of the NER-resistant CPD that showed slight over-twisting of −6 ± 2° (**Figure 4B**).

Next, the helix bend angle and the bend directions were analyzed for the same sequence region as in the unbound DNA simulations. Accompanying the untwisting upon Rad4 initial binding, the CCC/GGG DNA showed a larger bend angle (24 ± 5°) than the CGC/GCG case (12 ± 2°); the latter showed essentially no change in the bend angle upon Rad4 initial binding, while the CCC/GGG bend angle increased notably upon Rad4 initial binding, being 13 ± 1° in unbound DNA (**Figures S6B and S8B**). Another feature of the CCC/GGG DNA is that its bend becomes further directed towards the ‘open’ structure direction, with a bend direction pseudo-dihedral angle of −53 ± 7°, compared to −45 ± 8° for the unbound DNA. The bend direction of the CGC/GCG DNA (−17 ± 12°) is similar to that of the unbound DNA (−19 ± 10°) and oscillates greatly as it fails to be bound by the β−hairpin2, described below (**Figures S6C and S8C**).

In accordance with these structural differences in the two DNA duplexes, the CCC/GGG duplex exhibited weakened stacking for the 6-mer sequence, with van der Waals stacking interactions of -77.3 ± 1.3 kcal/mol, 3 kcal/mol higher than its unbound state (−80.3 ± 0.3 kcal/mol, **Figures S6D and S8D**). However, the CGC/GCG duplex had van der Waals stacking interactions of −81.5 ± 0.4 kcal/mol, unchanged from its unbound state (−81.6 ± 0.2 kcal/mol, **Figures S6D and S8D**).

Above all, the most conspicuous difference was in the way BHD2 engaged with the DNA. The aforementioned differences in the DNA bending and stacking energies could primarily arise from such differences in BHD2 binding. With CCC/GGG, the β-hairpin2 engaged with the DNA’s minor groove and the interaction was sustained. In contrast, the β-hairpin2 not only failed to engage with but also was expelled from the minor groove of CGC/GCG. These differences were further quantified by the BHD2-occupied alpha space (AS) volume (31), detailed in SI Methods. The computed AS volume reflects the curvature and surface area of the DNA minor groove occupied by BHD2. The AS volume was 165 (Å^3^) with the CCC/GGG sequence and 0 (Å^3^) with the CGC/GCG sequence. For comparison, the AS volumes with Rad4 were 349 (Å^3^) for 6-4PP and 110 (Å^3^) for CPD, respectively (**Figure 4B**) (7). The representative structures also showed that β-hairpin2 formed four hydrogen bonds with the DNA backbones of CCC/GGG but only two with those of CGC/GCG (**Figure 4A**). We remark that the free CCC/GGG DNA had already exhibited significant untwisting and slide in a direction that was conducive to accommodating the incoming β-hairpin2 in the minor groove whereas CGC/GCG lacked such intrinsic structural features. Thus, again, such results indicate that the inherent structural properties of CCC/GGG likely promote the ‘opening’ by Rad4 shown in the crystal structures. For CGC/GCG DNA, the exclusion of BHD2 from the minor groove led BHD2 to reside partially in the major groove (**Figure 4A**), which could further hinder the insertion of the β−hairpin3 into the DNA major groove to form an ‘open’ structure (**Figure S8E**). These MD simulations are thus consistent with the crystal structures.

## DISCUSSION

Previous studies have implicated various factors in the lesion recognition step of NER, including DNA conformations, lesion topology, stereochemistry, nature of the adducted base and sequence context (32–43). For many lesions, the more destabilizing and distorting a lesion is, the better it is recognized and repaired, and the sequence contexts that destabilize the lesion site could aid the recognition and repair efficiency of NER lesions. For instance, the 6-4PP UV-lesion is more destabilizing, distorting and dynamic than CPD when present in a duplex DNA. Accordingly, 6-4PP is much more efficiently recognized and repaired by XPC and NER than CPD (16,44–49). However, the recognition and repair rates of thymine-thymine-linked CPD improve dramatically if the T-T CPD is placed within a string of multiple mismatches against dT’s instead of with matched dA partners (6,17,18,50). We have also previously shown that different mismatch sequences have varying Rad4-binding specificities, which also correlated with the extent of the distortions induced by the mismatches. For instance, the more distorted and conformationally heterogeneous CCC/CCC mismatch had a greater specificity than a TAT/TAT mismatch that was structurally less distorted and heterogeneous. Furthermore, the more distorted sequences revealed more destabilization in thermal melting (51). However, it should be noted that thermal melting of a lesion-containing DNA duplex does not always correlate with NER. This has been discussed in (45), where it was explained that thermal stability is measured for a duplex of a given length while the opening needed for recognition involves the destabilization of a local segment.

The DNA in our study is matched DNA with consecutive or alternating C/G base pairs. The melting temperatures (Tm) of 24-bp DNA containing the CCC/GGG or CGC/GCG sequences were comparable to each other (76.5 ± 1.0 vs. 74.3 ± 1.8 °C) (**Figure S1 and S2**). However, the structures of the Δβ-hairpin3 mutant Rad4-tethered to these DNAs were dramatically different as discussed above: CCC/GGG DNA was able to form a ‘open-like’ conformation, while CGC/GCG DNA could not be opened but led the protein to bind to the ends of the DNA. These results suggest that (1) certain DNA sequences are more prone to being ‘opened’ than others; (2) for certain DNAs under tethered condition, binding to the DNA ends may be preferred over untwisting and opening; and (3) this preference does not necessarily correlate with the thermal stability of the DNA duplex as measured by T_m_.

Interestingly, our MD simulations with DNA in the absence of protein showed that CCC/GGG had higher intrinsic slide, roll and untwist and weaker base stacking energy. Furthermore, CCC/GGG was bent in a direction that is consistent with the direction of DNA in the ‘open’ structure. It has been shown that runs of guanines induce significant sliding due to steric hindrance caused by amino groups in adjacent guanines (28). Upon the initial binding with Rad4, BHD2-induced untwisting led to the approach of β-hairpin3 to the major groove side for further opening of the CCC/GGG sequence (**Figures S8A and S8E**), revealing the cooperative interplay between BHD2 and BHD3. Rad4 could take advantage of and amplify the intrinsically higher distortion/distort-ability and weaker van der Waals stacking energy in the CCC/GGG sequence. In general, the untwisting of the DNA duplex and the engagement of BHD2 shown with the CCC/GGG sequence recapitulates the features previously shown with NER-proficient lesions (7, 30). However, unlike the *bona fide* NER lesions such as 6-4PP or dibenzo[*a,l*]pyrene-derived dG lesion, we did not observe partner base flipping for CCC/GGG, which indicates that the kinetics of DNA opening for the more stable, matched DNA must be slower than those of the 6-4PP lesion. On the other hand, the stronger stacking for the CGC/GCG duplex supported the stability of its DNA even in the presence of WT Rad4, consistent with its failure to untwist and engage with the BHD2 hairpin in the minor groove, features which were also characteristic of NER-impaired/resistant lesions such as CPD (7, 30). As a result of these localized structural and energetic properties in the CGC/GCG duplex, the potential ‘open’ site is not sensed nor further processed by the BHD2 and BHD3 domains, thus congruent with its reverse-mode crystal structure showing ‘closed’ DNA.

### Implications of C/G-rich sequences

Several studies have implicated Rad4-Rad23 and XPC complexes in roles outside of the NER repair function, in particular, transcription. Furthermore, many of the gene regulatory DNA sequences reported to associate with Rad4/XPC possess a GGG-containing consensus sequence. In the study by Reed and colleagues, Rad4-Rad23 was shown to associate with STRE (Stress Response Element) promoter in the absence of UV light to regulate the transcription of several DNA damage repair signaling genes in yeast (52). The STRE elements are present in the upstream region of many genes, induced under various stress conditions such as osmotic pressure, oxidative stress and heat and contain a run of G’s (AGGGG)(53, 54). Also, previously, Tjian and colleagues reported that XPC serves as a stem cell coactivator required for OCT4/SOX2 transcriptional activation. Intriguingly, the consensus sequences of XPC/RAD23B colocalization includes KLF4 (nCCnCnCCCn) and SP1 (CCCCnCCCCC) (21, 22) that are also enriched with strings of G’s. Le May and colleagues recently reported that XPC colocalizes with RNA polymerase II in the absence of damage and functions as a co-activator for recruiting the ATAC transcription coactivator complex to promoters by interacting with E2F1(55). Interestingly, the E2F1 consensus sequence contains runs of G’s (NNGGCGGGAA, http://homer.ucsd.edu/homer/motif/HomerMotifDB/homerResults/E2F1.html).

In light of our current findings, the sequences containing runs of G’s may be more susceptible to Rad4/XPC-induced DNA opening and thus more likely to be specifically recognized than other sequences in the absence of DNA damage but in the context of a situation where prolonged residence time is allowed. We therefore envision that our findings may also help decipher the role of XPC in processes other than DNA repair.

## Acknowledgements

We thank the staff of Advanced Photon Source LS-CAT beamline for the help with data collection. We also thank the members of the Min and Broyde groups for their support.

---

**Table S1.**
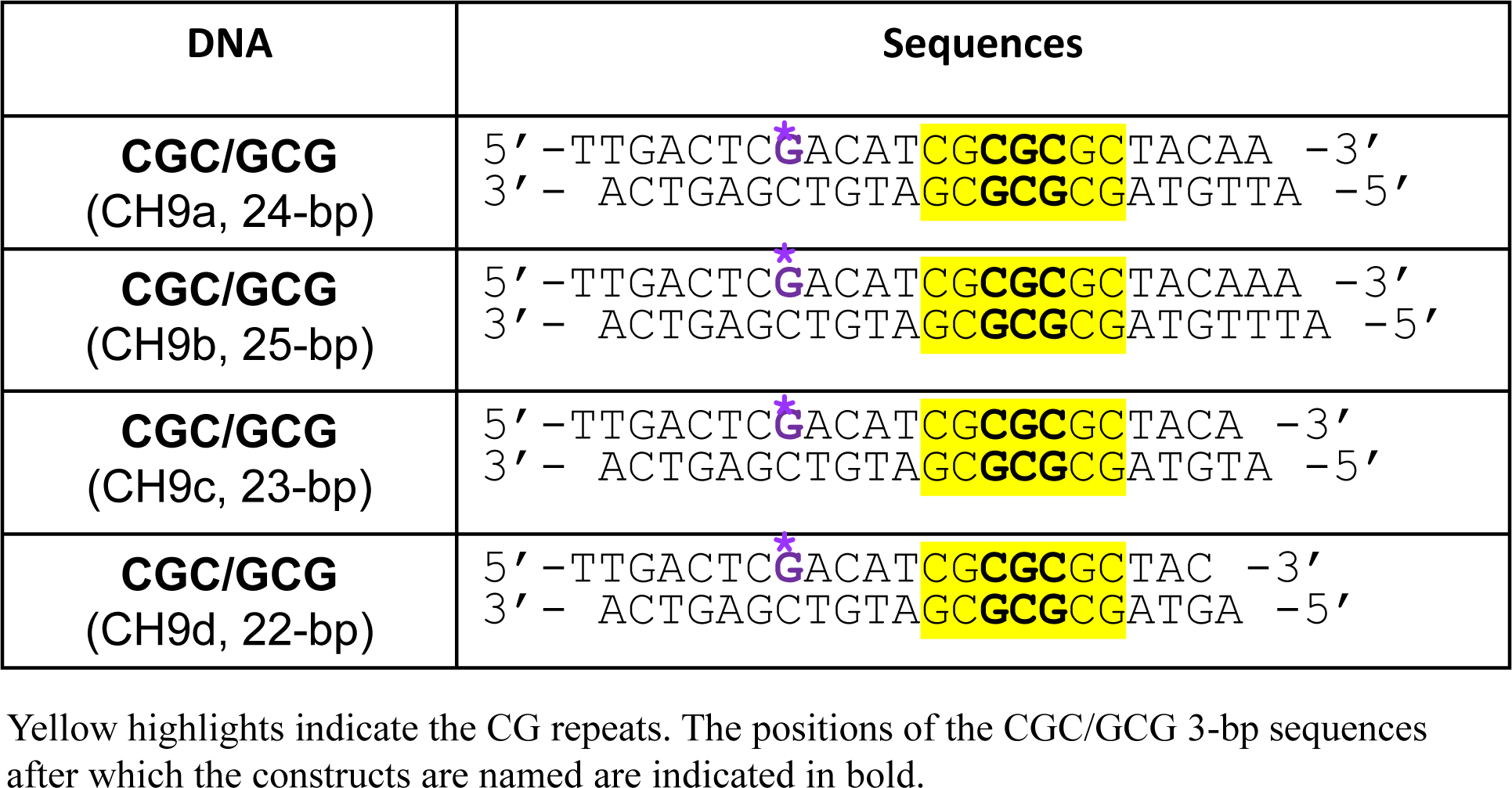
Sequences of crosslinkable DNA constructs containing G* in purple used for the crystallization trials.

| DNA | Sequences |
| --- | --- |
| **CGC/GCG**<br>(CH9a, 24-bp) | 5' - TTGACTC<sup>*</sup>GACAT**CGCGCGC**TACAA -3'<br>3' - ACTGAGCTGTA**GCGCGCG**ATGTTA -5' |
| **CGC/GCG**<br>(CH9b, 25-bp) | 5' - TTGACTC<sup>*</sup>GACAT**CGCGCGC**TACAAA -3'<br>3' - ACTGAGCTGTA**GCGCGCG**ATGTTTA -5' |
| **CGC/GCG**<br>(CH9c, 23-bp) | 5' - TTGACTC<sup>*</sup>GACAT**CGCGCGC**TACA -3'<br>3' - ACTGAGCTGTA**GCGCGCG**ATGTA -5' |
| **CGC/GCG**<br>(CH9d, 22-bp) | 5' - TTGACTC<sup>*</sup>GACAT**CGCGCGC**TAC -3'<br>3' - ACTGAGCTGTA**GCGCGCG**ATGA -5' |

Yellow highlights indicate the CG repeats. The positions of the CGC/GCG 3-bp sequences after which the constructs are named are indicated in bold.

**Table S2.** Data collection and refinement statistics (molecular replacement)

|  | 6UG1 (SC41bxCH9c) |
| --- | --- |
| **Data collection** |  |
| Wavelength (Å) | 0.9794 |
| Space group | P 1 |
| Cell dimensions |  |
| *a*, *b*, *c* (Å) | 53.2 59.6 78.2 |
| $\alpha$, $\beta$, $\gamma$ (°) | 105.5 97.9 107.1 |
| Resolution (Å) | 50.0 – 2.9 |
| *R*<sub>sym</sub> or *R*<sub>merge</sub> | 11.6 (51.1) |
| *I* / $\sigma I$ | 9.7 (2.2) |
| Completeness (%) | 90.8% (74.1%) |
| Redundancy | 3.4 (2.9) |
| **Refinement** |  |
| Resolution (Å) | 37.4-2.8 (2.93-2.83) |
| No. reflections | 17956 (1519) |
| *R*<sub>work</sub> / *R*<sub>free</sub>(%) | 22.57/27.39 |
| No. atoms | 5231 |
| Protein | 4300 |
| DNA | 931 |
| Water | 0 |
| *B*-factors (Å<sup>2</sup>) | 76.8 |
| Protein | 62.3 |
| DNA (G47) | 91.3(91.7) |
| Water | 0 |
| R.m.s. deviations |  |
| Bond lengths (Å) | 0.005 |
| Bond angles (°) | 0.97 |

R.m.s., root mean squared.
\*Values in parentheses are for highest-resolution shell.

## SI Method

### Streak-seeding

To obtain larger crystals, the crystal showers were used for streak-seeding by transferring 5–10 crystals (∼20 μm) into a 10 µl drop of harvest buffer (50 mM BTP-HCl, 200 mM NaCl and 12% isopropanol pH 6.8). The crystals were smashed using a fine needle and were transferred into a tube containing 250 µl of the harvest buffer, which was then mixed by gentle vortexing. The freshly prepared seed-stocks always produced better quality crystals. The best diffracting crystal were obtained from drops pre-equilibrated for 1 h (50 mM BTP-HCl, 200 mM NaCl and 0-5% isopropanol, pH 6.8), seeded by passing the tip of a cat’s whisker dipped into a fresh seed-stock solution through the drop. The crystals grew to maximum size of ∼70-80 μm in 10-12 days.

### Melting temperatures of DNA duplexes used in this study

The overall thermal stabilities of the DNA duplexes (**Figures S1 & S2**) were measured as follows. The absorbance at 260 nm of each DNA duplex (1.5 μM) was measured in a sample cuvette of path length 1 cm, using a Cary 300 Bio UV-Visible spectrophotometer equipped with a Varian temperature controller. The absorbance measurements were done from 30 to 85 °C at every 1.0 °C interval. The absorbance data were subsequently averaged (smoothened) over a 2.5°C (5 data points) window prior to calculating derivatives using Origin software OriginPro. Melting temperatures were then determined from the melting profiles using methods described previously (13, 18).

### Initial models, water box and counterions

The initial models of the two 13-mer DNA sequences (nucleotide steps 10 – 22, Figure 2 in the main text) were created using standard B-DNA in Discovery Studio 2.5 (Accelrys Software Inc.). The DNA models were neutralized with Na^+^ counterions and solvated with explicit TIP3P water(1) in a cubic periodic box with side length of 59.0 Å using the tLEAP module of the AMBER16 suite of programs (2). The initial models of the Rad4 docked to the two DNA sequences were prepared as in earlier work (3), where the BHD2 and BHD3 domains are close but not bound to the sequences yet. The protein-DNA complexes were neutralized with Na+ counterions and solvated with explicit TIP3P water (1) in a cubic periodic box with side length of 125.0 Å using the tLEAP module of the AMBER16 suite of programs (2).

### MD simulations

We used the ff14SB force field for the MD simulations (4). All MD simulations were carried out using the AMBER16 suite of programs (2). The Particle-Mesh Ewald method (5) with 9.0 Å cutoff for the non-bonded interactions was used in the energy minimizations and MD simulations. Minimizations were carried out in three stages. First, 500 steps of steepest descent minimization followed by 500 cycles of conjugate gradient minimization were conducted for the water molecules and counterions with a restraint force constant of 50 kcal/(mol·Å^2^) on the solute molecules. Then, 500 steps of steepest descent minimization followed by 500 cycles of conjugate gradient minimization were carried out for the water molecules and counterions with a restraint force constant of 10 kcal/(mol·Å^2^) on the solute molecules. In the last round, 500 steps of steepest descent minimization followed by 500 cycles of conjugate gradient minimization were carried out on the whole system without restraints. The minimized structures were then subjected to three rounds of equilibration. First, each system was equilibrated at constant temperature of 10° K for 30 ps with the solute molecules fixed with a restraint force constant of 50 kcal/(mol·Å^2^). Then the system was heated from 10° K to 300° K over 300 ps with the solute molecules fixed with a restraint force constant of 50 kcal/(mol·Å^2^) at constant volume. In the last round of equilibration, the restraint force constant on the solute was reduced through three steps: at 10 kcal/(mol·Å^2^) for 200 ps, at 1 kcal/(mol·Å^2^) for 200 ps, and then at 0.1 kcal/(mol·Å^2^) for 200 ps with constant pressure at 300° K. Following equilibration, production MD simulations for each system were carried out in a constant-temperature, constant-pressure (NPT) ensemble at 300° K and constant pressure of 1 Atm for 2 μs. The temperature was controlled with a Langevin thermostat (6) with a 5 ps^-1^ collision frequency. The pressure was maintained with the Berendsen coupling method (7). A 2.0 fs time step and the SHAKE algorithm (8) were applied in all MD simulations.

### Structural analyses

#### Principle component analysis (PCA) and RMSD analysis

PCA analyses for the ensemble of structures from MD simulations of the DNA duplexes were performed with heavy atoms of the DNA sequences excluding flexible end base pairs, using R (9) with the Bio3D package (10). The structures from each ensemble were superposed to the first frame at the heavy atoms of the base pairs 3’ to the run of guanines and serve as our reference for structural changes due to the sequence effect. Due to the fact that each ensemble provides only one cluster, no clustering was performed. The RMSD for the heavy atoms of the DNA sequences excluding flexible end base pairs were calculated for the ensemble of structures in the production MD, using the cpptraj module of AMBER 16 (2). PCA analyses for the Rad4 initial binding to the DNA duplexes were carried out as described in earlier work (11). The RMSD for the heavy atoms of the BHD2 domain and potential open site 6-mer DNA sequences (nucleotide steps 14 – 19, **Figure 2** in the main text) were calculated for the ensemble of structures in the production MD. The best representative structure for each ensemble is defined as the one frame that has the shortest RMSD for the heavy atoms used in the calculation of RMSD values to all other frames.

#### Base pair step parameters

We measured base pair step parameters (shift, tilt, slide, roll, rise and twist, **Figure S6** and **Figure 3** in the main text) between each base pair step of the 13-mer duplexes excluding the flexible end base pairs, using the cpptraj module of AMBER16 (2).

#### Helix bend and DNA tail bending direction pseudo-dihedral angle

We measured the total helix bend angle for the duplex excluding flexible end base pairs using Curves+ (12) (**Figure S6**). The DNA tail bend direction was measured using a pseudo-dihedral angle, defined in **Figure S6**, using the cpptraj module of AMBER16 (2).

#### Van der Waals stacking interaction energy

The van der Waals stacking interaction energy for the base pairs of the potential open site 6-mer sequences (base pair steps 14 – 19 in **Figure 2** of the main text) were calculated using the cpptraj module of AMBER16 for the Lennard-Jones potential(2).

#### Analysis of DNA Untwisting upon Rad4 binding

The untwisting of the lesion-containing DNA along the initial binding trajectory was monitored by the Untwist angle defined as Untwist = Twist initial –Twist. The twist angle was measured between the end base pairs of the lesion-containing 6-mer (base pair steps 14 and 19 in **Figure 2** of the main text) using the cpptraj module of AMBER16(2). Twist initial is the ensemble average twist angle of the lesion-containing 6-mer during the first 1 ns of production MD during which significant untwisting was not observed (**Figure S8**); this ensemble represents the state of the lesion-containing sequence before the engagement of Rad4, especially BHD2. Positive values indicate further untwisting and negative values indicate further twisting.

#### The AlphaSpace volumes analyses for BHD2 binding to the minor groove

The best representative structure for the initial binding state of each lesion-containing duplex was further analyzed to quantify BHD2’s binding into the DNA minor groove around the lesion site. The alpha space volumes of the binding pockets in the DNA and their occupancies by BHD2 were calculated using AlphaSpace v1.0(13): the lesion-containing DNA was set as the receptor and BHD2 was set as the ligand. The total occupied alpha space volume was used to quantify the extent of BHD2 binding into the DNA minor groove (**Figure S8**). The value reflects the curvature and surface area of the DNA minor groove region that is occupied by BHD2.

#### Analysis of hydrogen bonding between BHD2 and DNA duplexes

Hydrogen bonds between BHD2 and the DNA duplexes were counted for the best representative structures in each ensemble using each pair of donor and acceptor atoms that has a hydrogen bond (heavy-light-heavy atom) angle ≥145° and heavy-to-heavy atom distance ≤ 3.3 Å.

#### Block average analyses

The DNA structural parameters and van der Waals energies for the stable ensemble of each MD simulation were analyzed using the block averaging method (14, 15). In brief, the time series data were divided into “blocks” with a block size that exceeds the longest correlation time, 50 ns in our case. The average for each block was computed and termed “block average”. The mean values and the standard deviations of the block averages given in the main text were used to represent the average and the variance of averages.

Molecular structures were rendered using PyMOL 1.3.x (Schrodinger, LLC.) or UCSF Chimera (developed by the Resource for Biocomputing, Visualization, and Informatics at the University of California, San Francisco, with support from NIH P41-GM103311) (16). All MD simulation data were plotted using MATLAB 7.10.0 (The MathWorks, Inc.).

## SI DISCUSSION

As described in the main text, the DNA in this ‘reverse mode’ structure was obtained with a 23- bp matched DNA containing CGC/GCG sequence whereas the ‘open’ or pseudo-open structures solved so far contained 24-bp DNA with various lesions and sequences. Here, we argue that the differences in the DNA lengths did not likely cause the different binding modes but rather played a role in capturing existing conformation(s) in solution into crystals by promoting crystal packings for a given conformation.

First of all, it can be structurally demonstrated that adding or removing one nucleotide from one of the DNA ends (L-side) in a 23-bp or 24-bp DNA would not impact any known protein-DNA contacts in either binding modes. For instance, if the protein were to bind in a ‘reverse mode’ and cap the end of a 24-bp DNA (for instance, containing CCC/GGG), it should have been allowed just the same as was seen in the 23-bp CGC/GCG DNA, as the S-side sequences of the DNA of both DNA are the same. Conversely, if the protein were to ‘open’ the CGC/GCG DNA, the 23-bp construct had a length sufficient for making all the contacts with the protein in an ‘open’ conformation, as the contacts on L-side of the DNA does not extend much beyond the potential ‘open’ site where nucleotides get flipped out which is present in both the 23- and 24-bp DNA(11). However, such alternative structures have not been observed and the crystals of Rad4 tethered to a 23-bp CCC/GGG or with 24-bp CGC/GCG did not diffract well (data not shown). This indicates that these lengths of DNA were not likely optimal for the crystallographic packing of the given complex structure. In other words, the crystals we obtained with 23-bp CCC/GGG were not likely to be in ‘reverse mode’ (which would have preferred 23- bp for packing) and the crystals we obtained with 24-bp CGC/GCG were not likely to be in an ‘open’ structure (which would have preferred 24-bp). Thus, we propose that the differences in the ‘reverse’ vs ‘open(-like)’ mode of binding for CGC/GCG and CCC/GGG sequences are therefore not likely instigated by the 1-bp difference in the lengths but by the other difference in the sample which is the differences in the central DNA sequences. (We note, however, that we cannot entirely exclude the possibility that minor populations of alternative conformations could exist in solution that were not captured by the crystal structures.)

## SI Figure and Figure Legends

**Figure S1.**
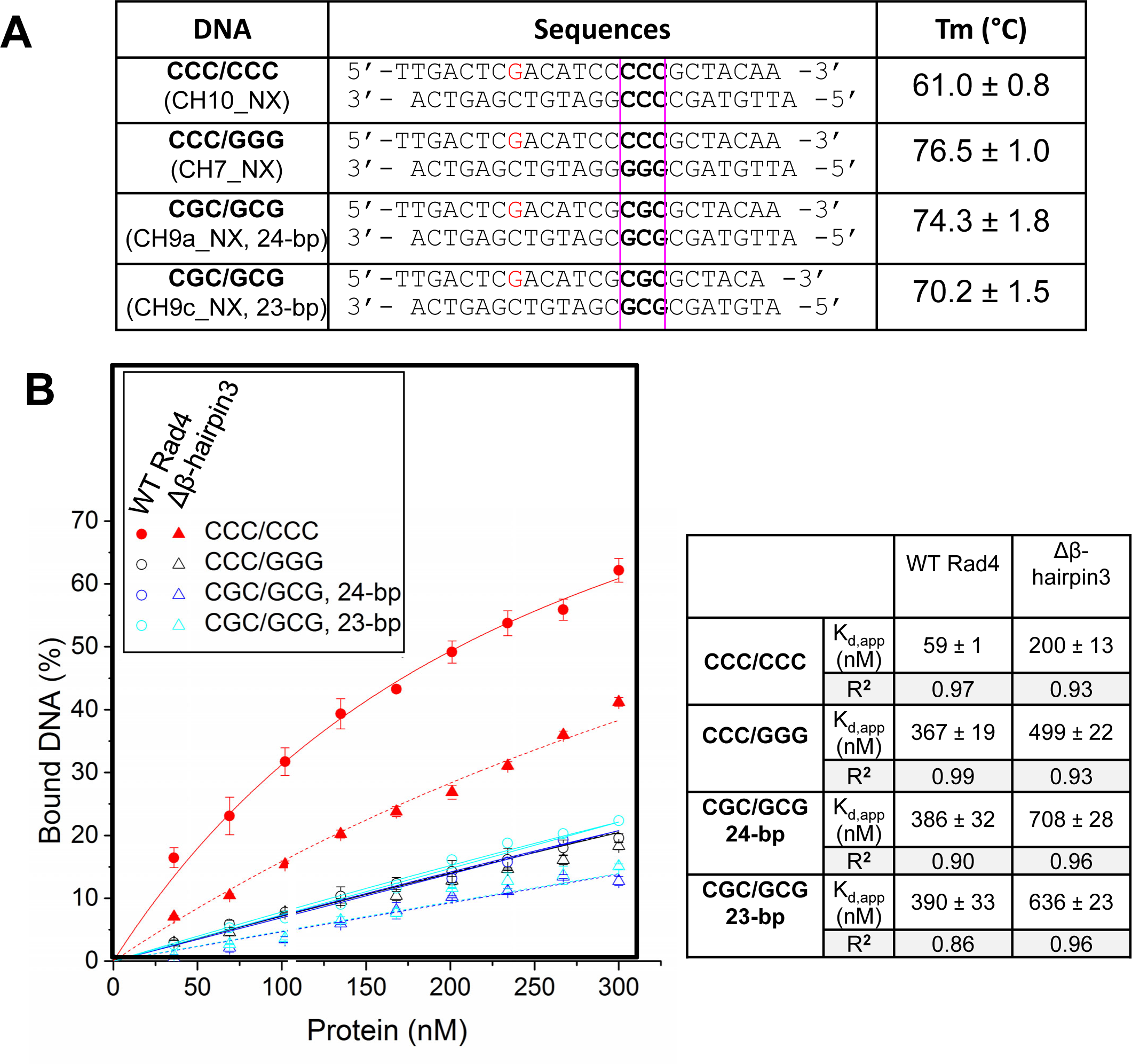
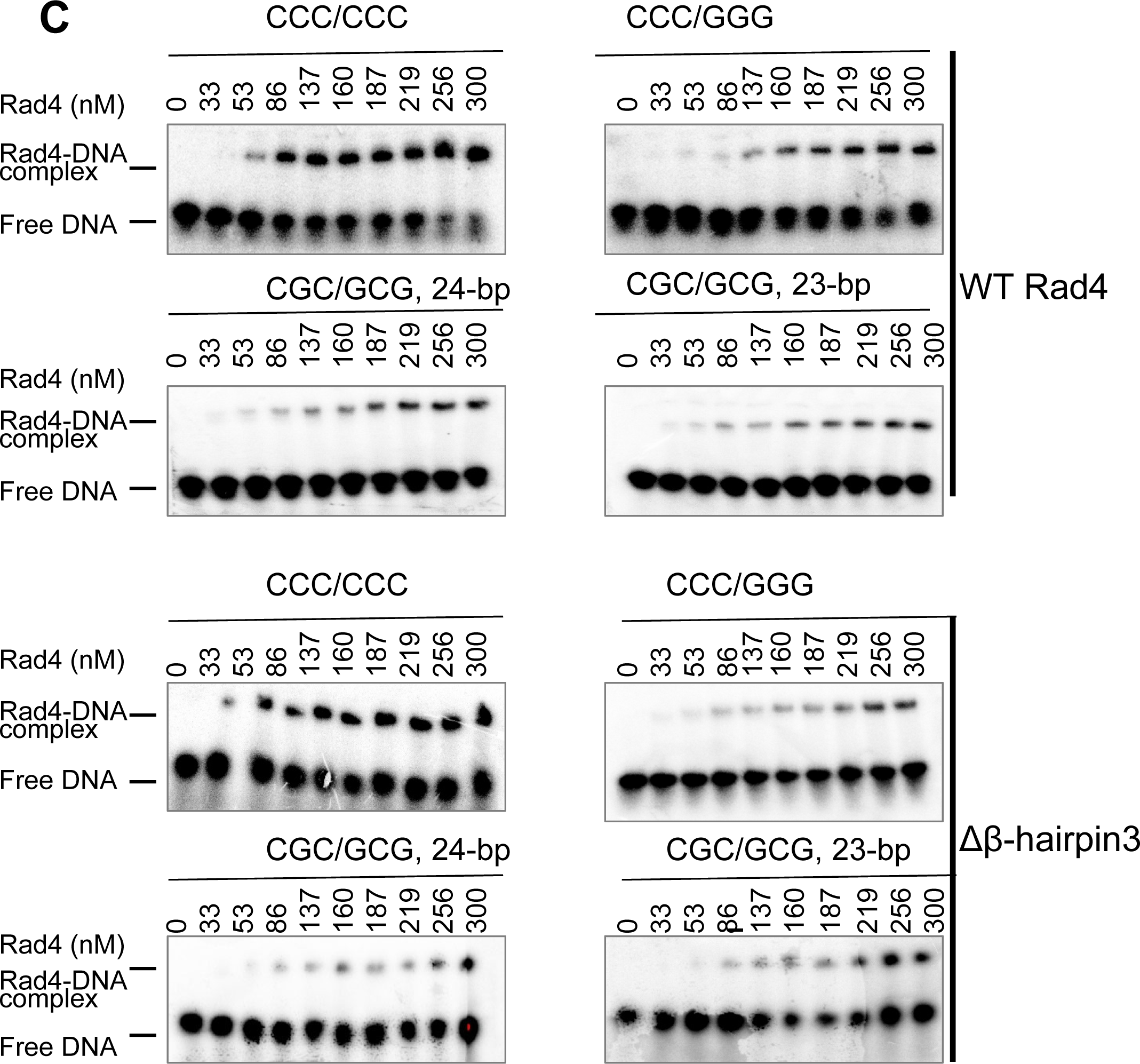
Characterization of the apparent binding affinities between Rad4 and DNA. **(A)** The construct names, sequences and melting temperatures of duplexed DNA used for the competitive EMSA assays. The positions of the 3-bp sequences after which the constructs are named (e.g., CCC/CCC in the CCC/CCC construct) are indicated in bold and boxed in pink. The deoxyguanosines marked red indicate the positions of the crosslinkable dG* used for crystallization studies (e.g., **Figure 1B**). **(B)** Quantification of the bound DNA fractions versus the concentrations of the protein from the EMSA. Symbols of different colors indicate different DNA sequences; filled and empty symbols indicate mismatched and matched DNA, respectively; circles and triangles indicate WT and Δβ-hairpin3, respectively. The error bars indicate ±standard deviations from triplicate gels. Solid and dotted lines indicate the fit curves of the data points for WT and Δβ-hairpin3, respectively. The apparent Kd values for specific binding to various DNAs were calculated as described (11,17,18). These results showed that Δβ-hairpin3 mutant had weakened specific binding but retained most affinities for nonspecific binding, resulting in a significant loss in the lesion recognition specificity. Neither the WT nor mutant protein showed significant difference in binding to 23 bp versus 24 bp nonspecific CGC/GCG DNA. **(C)** Representative gel images used for (B), showing the binding of different DNA sequences to WT (top) and Δβ-hairpin3 mutant Rad4 (bottom).

**Figure S2.**
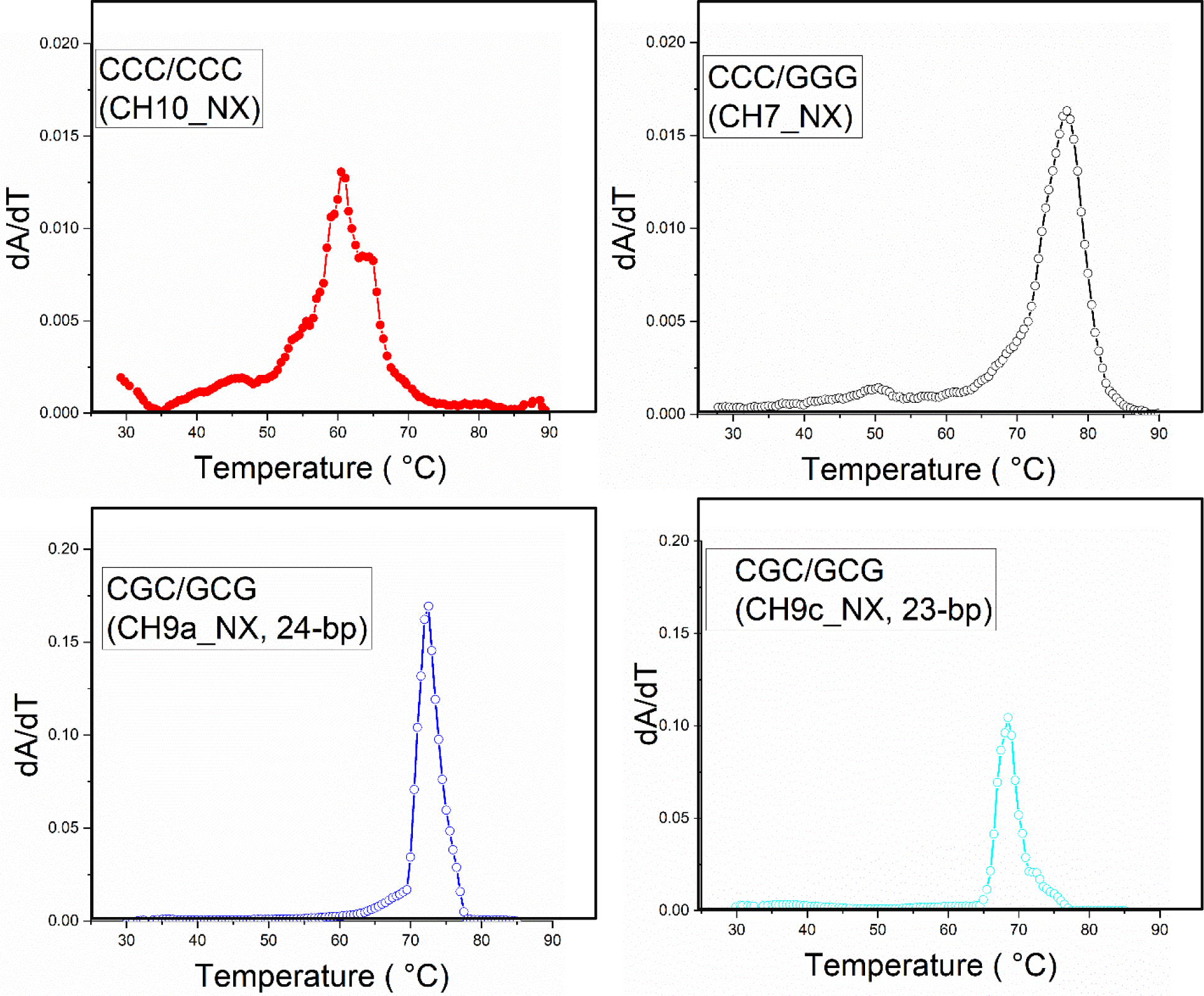
Melting temperature measurements of double-stranded DNA substrates. Melting temperature profiles for the Tm’s reported in **Figure S1**. See Figure S1 for the sequences of each construct. The color and symbol used for the different DNA sequences are the same as in Figure S1B.

**Figure S3.**
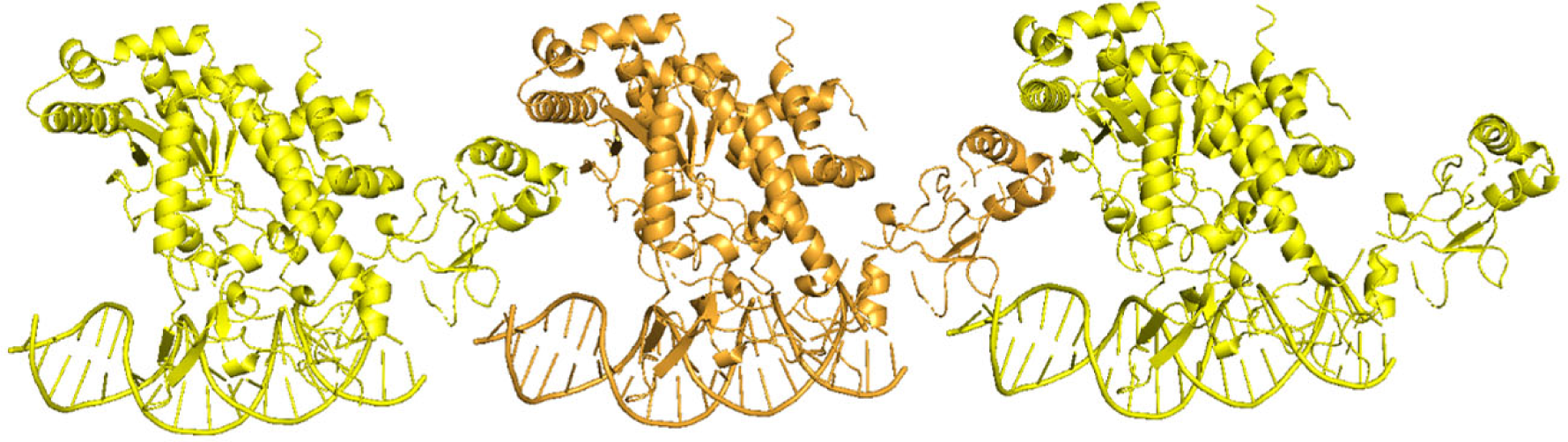
Crystal packing of the Δβ-hairpin3 tethered to CGC/GCG DNA. In this *P1* space group crystal, the BHD2/3 packs against the L-side of the neighboring DNA molecules to make the complexes arrange in a head-to-tail manner.

**Figure S4.**
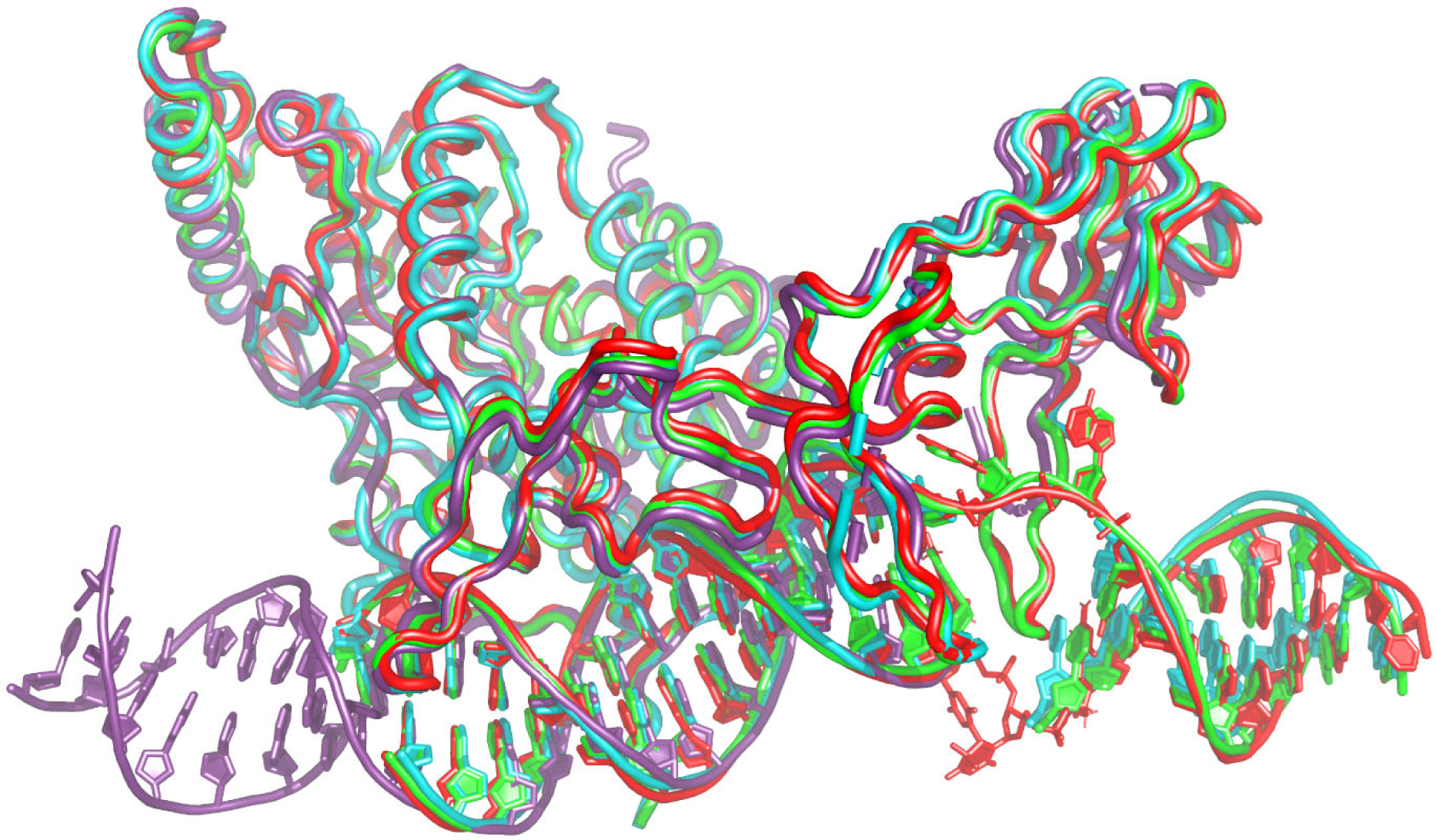
Superposition of the ‘reverse mode’ and ‘(pseudo-)open’ Rad4-DNA crystal structures. Δβ-hairpin3 crosslinked to CGC/GCG (PDB ID: 6UGI, purple) shows the ‘reverse mode’ binding. The WT Rad4 bound to 6-4PP (6CFI, red) and the WT crosslinked to CCC/GGG (4YIR, green) both show ‘open’ conformation; Δβ-hairpin3 crosslinked to CCC/GGG (6UBF, cyan) shows ‘pseudo-open’ structure.

**Figure S5.**
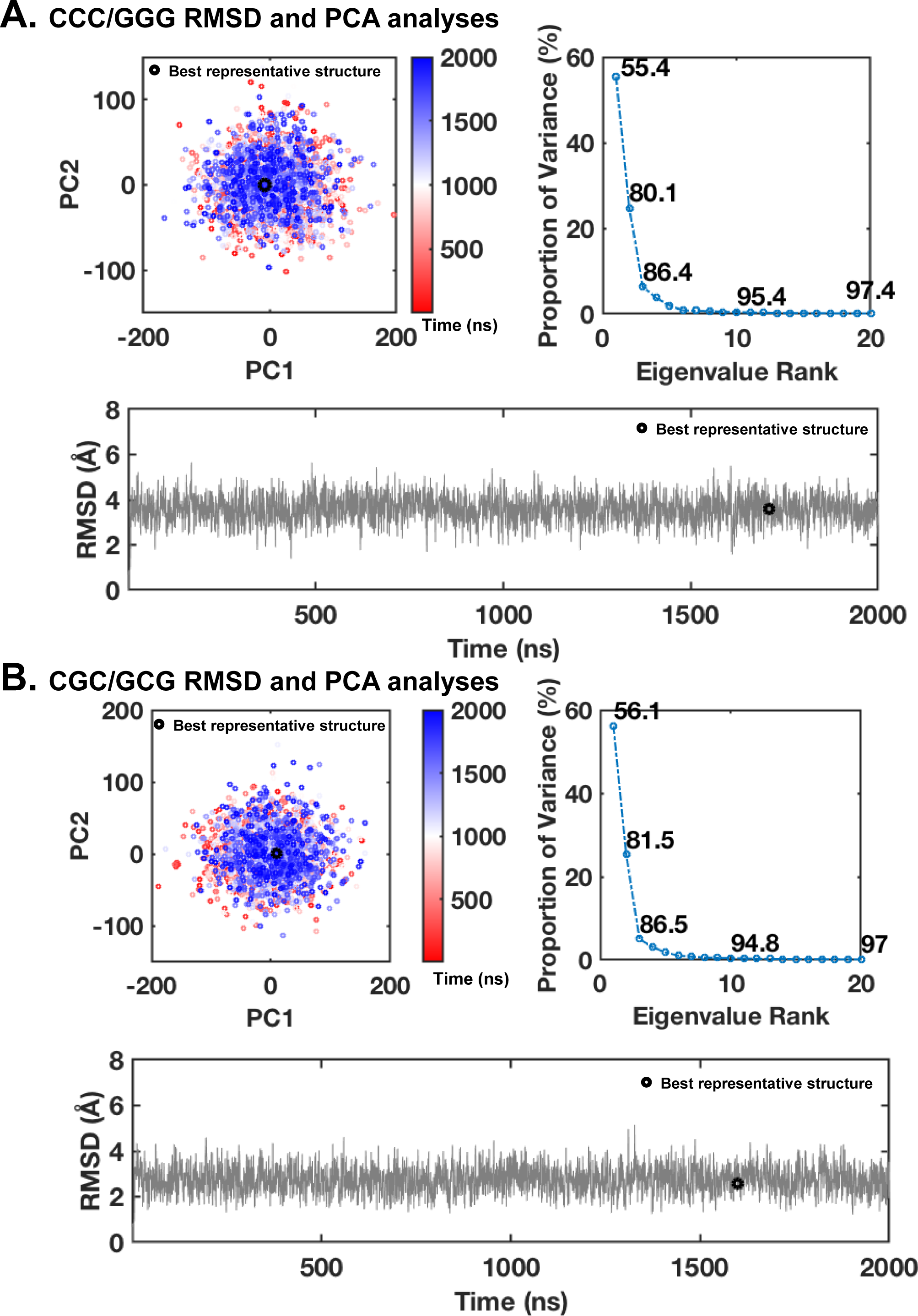
Principle component analyses and RMSD for the MD simulations of the (A) CCC/GGG and (B) CGC/GCG duplexes. The 0 to 2 μs ensemble was used for further analyses, since all parameters were stable (single cluster in the PCA plot and stable RMSD values).

**Figure S6.**
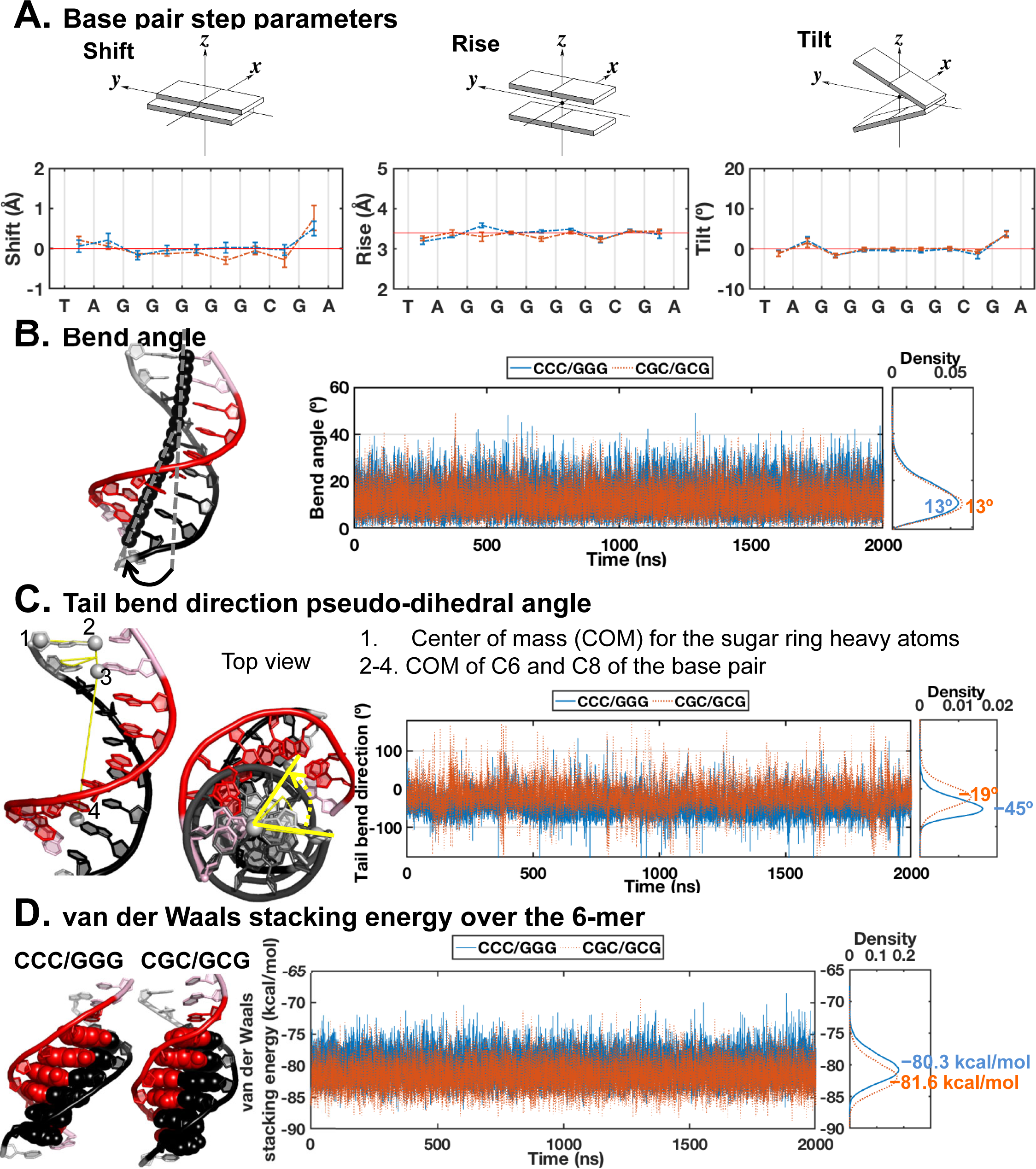
Structural analyses for the MD simulations of the unbound CCC/GGG and CGC/GCG duplexes. **(A)** Shift, rise and tilt base pair step parameters. The block averaged means and standard deviations for the parameter values are shown. Illustrations of the base pair step parameters are adapted from 3DNA (19) **(B)** Bend angles. The best representative structure of CCC/GGG is shown on the left to illustrate the bend angle. The helix axis is shown in black spheres. **(C)** DNA tail bend direction pseudo-dihedral angles. The best representative structure of CCC/GGG is shown on the left to illustrate the definition of the tail bend direction pseudo-dihedral angle, with the centers of mass (COM) utilized in the computation shown as white spheres. The top view shows the tail bend direction compared to B-DNA in dark gray. **(D)** van der Waals stacking energy for the potential open site 6-mer (base pair steps 14 – 19 in Figure 2). The best representative structures of CCC/GGG and CGC/GCG are shown on the left with the base pairs of the 6-mer used for the calculation of the van der Waals energy in spheres. Time dependent values are shown for properties in panels (B), (C) and (D). Kernel densities are calculated for the raw data using the ksdensity function with 200 bins in MATLAB 7.10.0 (The MathWorks, Inc.), and are plotted on the side with mean values labeled. These kernel densities are representative of the population distributions over the range of each property (e.g., bend angle).

**Figure S7.**
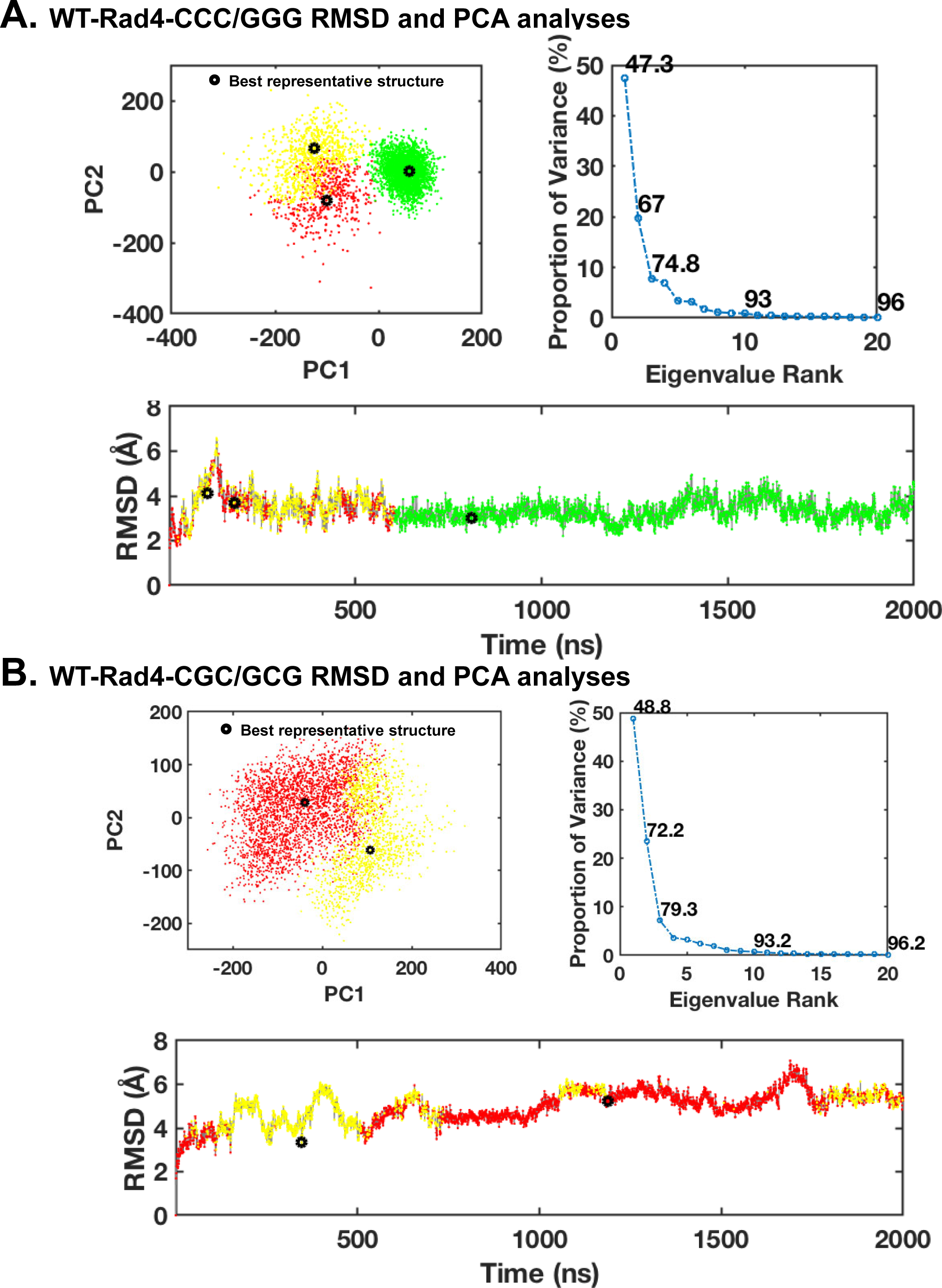
Principle component analyses and RMSD for the MD simulations of the Rad4 initial binding to the (A) CCC/GGG and (B) CGC/GCG duplexes. The 1 to 2 μs ensemble was used for further analyses, since one main structure cluster is achieved in ∼ 1 μs (with stable RMSD value). Different structure clusters are color-coded. The best representative structure values for each cluster are shown in black circles.

**Figure S8.**
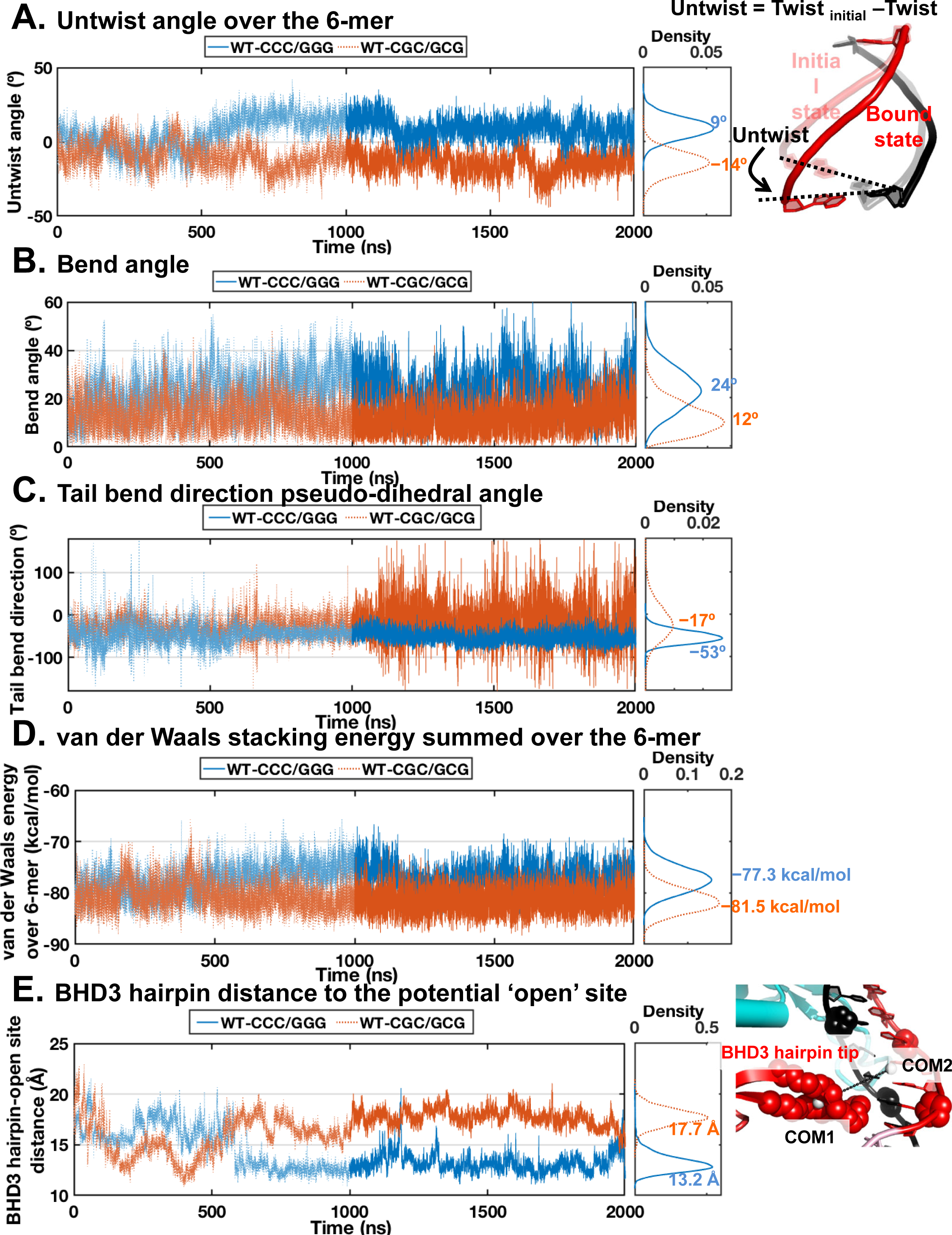
Structural and energetic analyses for the MD simulations of the Rad4 initial binding to the CCC/GGG and CGC/GCG duplexes. **(A)** Untwist angles illustrated on the right, **(B)** bend angles, **(C)** DNA tail bend direction pseudo-dihedral angles, **(D)** van der Waals stacking energy for the potential ‘open’ site 6-mer (base pair steps 14–19 in **Figure 2**), and **(E)** the distance between β−hairpin3 and the center of the potential ‘open’ site. The distance is calculated between the center of mass (COM) of the β−hairpin3 (residues 597-607) backbone heavy atoms (COM1) and the COM of sugar ring heavy atoms of the nucleotides surrounding the open site (nucleotide steps 15 and 18 in **Figure 2**) (COM2), illustrated on the right. Time dependent values are shown for all properties, with the values for initial binding states (1 – 2 μs) in darker shade. Kernel densities for the values of the initial binding states are calculated using the ksdensity function with 200 bins in MATLAB 7.10.0 (The MathWorks, Inc.), and are plotted on the side with mean values labeled. These kernel densities are representative of the population distributions over the range of each property (e.g. untwist angle).

